# Beta-caryophyllene restores blood pressure, heart rate, and electrocardiographic alterations while mitigating cardiometabolic biomarkers in a rat model of metabolic syndrome: Implications for non-communicable disease prevention

**DOI:** 10.64898/2026.09.09.750314

**Authors:** JolaOluwa Oluwatosin Yesufu, Benson Ekunola Olufemi, Abiodun Adegoke Adeyemi, Adesoji Adedipe Fasanmade

**Affiliations:** Department of Epidemiology and Medical Statistics, Faculty of Public Health, College of Medicine, University of Ibadan; Department of Veterinary Medicine, University of Ibadan, Ibadan.; Department of Pharmacognosy and Herbal Medicine, Faculty of Pharmacy, University of Ibadan; Department of Medicine, Faculty of Clinical Sciences, College of Medicine, University College Hospital, Ibadan/University of Ibadan.; Department of Physiology, Faculty of Basic Medical Sciences, College of Medicine, University of Ibadan.

**Keywords:** Cardiovascular dysfunction, Metabolic Syndrome, Beta-caryophyllene, High-fat-high-fructose diet, Public Health Nutrition, NCD Prevention

## Abstract

Metabolic Syndrome (MS) is an escalating non-communicable disease (NCD) and public health crisis heavily driven by modern high-calorie dietary transitions. MS is closely linked to catastrophic cardiovascular dysfunction (CVD). Conventional CVD medications are frequently constrained by high financial costs, system-wide adverse effects, and poor population-level accessibility in lower-middle-income countries (LMICs). This underscores an urgent public health need to evaluate safe, scalable, and low-cost dietary nutraceuticals for primary prevention. This study evaluated the cardioprotective and preventative effects of Beta-caryophyllene (BCP)— an affordable, widely accessible, and GRAS-designated dietary cannabinoid—in a lifestyle-mimicking rodent model of MS.

**Methods:** Seventy-two male Wistar rats (10 weeks old) were divided into six groups (n=12/group). Groups I and II served as controls given Normal Diet (ND) alone and ND+β-caryophyllene (50 mg/kg p.o.), respectively. Metabolic Syndrome was induced in Groups III–VI using a High-Fat High-Fructose Diet (HF/HFD) for 16 weeks. For the subsequent 4 weeks, Group III continued on HF/HFD, Group IV received HF/HFD + BCP (50 mg/kg p.o.), Group V underwent dietary withdrawal (returned to a normal diet [ND]), and Group VI received ND + BCP. Conscious hemodynamic variables and electrocardiograms (ECG) were recorded via telemetry. Plasma apolipoprotein-B (Apo-B) and C-reactive protein (CRP) were quantified using ELISA. Lipid profiles, oxidative stress parameters, and organ histopathology were evaluated using standard biochemical and staining methods, with data analysed using ANOVA at α = 0.05.

**Results:** In Group II, BCP decreased SBP (145.00±12.13 vs 120.30±2.15 mmHg), DBP (105.70±10.80 vs 94.00±17.20 mmHg), heart rate (448.00±11.90 vs 280.00±18.70 beats/min), and PR-interval (49.50±3.53 vs 45.00±12.70 ms) compared with Group I. Group IV significantly decreased SBP (129.00±1.28 vs 125.5±11.1 mmHg) and heart rate (380.00±16.20 vs 351.00±3.00 beats/min) compared with Group III. Furthermore, BCP significantly increased HDL-cholesterol (10.80±1.70 vs 15.40±1.00 mg/dl), SOD (0.35±0.07 vs 0.76±0.30 U/mg protein), and GSH (1.64±0.02 vs 3.60±0.13 mM) in Group IV compared with Group III. Conversely, BCP significantly decreased total cholesterol (166±31.00 vs 113±18.50 mg/dl), Apo-B (26.70±1.68 vs 19.00±1.30 mg/dl), leptin (5.10±0.03 vs 2.18±0.01 ng/ml), NO (6.30±0.40 vs 4.30±2.61 μM), and malondialdehyde (11.80±4.10 vs 9.90±2.30 μM) in Group IV. BCP significantly decreased triglycerides (64.90±1.30 vs 55.90±1.30 mg/dl) in Group VI compared with Group IV, while lowering CRP across all groups. Distortion of the cytoarchitecture of the heart, liver, and pancreas observed in Group III was completely reversed by BCP treatment in Group IV.

**Conclusion:** Beta-caryophyllene effectively ameliorates diet-induced metabolic syndrome and associated cardiovascular dysfunction via potent anti-atherogenic, anti-inflammatory, and antioxidant mechanisms. These findings suggest that integrating affordable, culturally accepted, BCP-rich indigenous functional ingredients into regional nutrition frameworks represents a highly scalable, low-cost public health strategy for global NCD prevention.

## Introduction

Despite being just a fist-sized organ, the human heart is the body’s strongest muscle^[1]^. Long before birth, often between 21 and 28 days after conception, the heart begins to beat in the uterus. Over a 70-year lifespan, the average heart beats roughly two and a half billion times, or 100,000 times every day^[1]^. The heart circulates blood throughout the body with each beat. During circulation, waste materials are eliminated by the liver. This extraordinary system can be attacked and weakened by a variety of events, many of which are curable and preventable^[2]^. Cardiovascular diseases (CVDs) remain the world’s largest cause of fatalities and permanent disabilities, accounting for approximately 17.9 million deaths per year^[3, 2]^. The CVDs include cardiac, rheumatic, and cerebrovascular disorders. Most CVD fatalities are caused by heart attacks and strokes, and 30% of them occur in those under 70^[3, 2.]^ In addition to being overweight or obese, people at risk for CVD may exhibit elevated blood pressure, glucose and cholesterol levels^[2]^. Despite the fact that many CVDs are avoidable, their prevalence is still rising, mostly due to insufficient preventive interventions^[2]^. Risk indicators for cardiovascular disease have evolved considerably across all demographic groups^[2]^. Tobacco, alcohol, obesity, elevated blood pressure and cholesterol concentrations are the five danger signs that account for at least thirty percent of all CVD in developed nations^[2]^.

Chronic diseases arise from pathological conditions that are irreversible, persistent, and lead to latent disability, and may require ongoing medical supervision and care^[4]^. The prevalence of metabolic illnesses is rising at an earlier age due to increased intake of fatty and sugar-rich foods and growing demand for them. Fructose is especially used in processed foods and fizzy drinks ^[5]^. Metabolic diseases such as diabetes mellitus, high blood pressure, hyperlipidemia and obesity, have shown altered cardiac autonomic function and metabolic syndrome is a constellation of these disorders^[6]^. This group of risk factors described as Metabolic Syndrome (MS) appears to promote the onset of chronic illness^[6]^. Metabolic Syndrome affects about 20 to 25 percent of adults worldwide. Those who have the Syndrome are threefold more likely to have a heart attack or stroke and twice as likely to die from it ^[7]^. A new CVD epidemic is largely driven by the Metabolic Syndrome’s constellation of heart disease indicators^[8]^. More than four out of ten people are unaware that they have diabetes, and 11.1%, or one in nine, of adults aged 20 to 79 have the disease^[8]^. Additionally, it has been predicted that by 2050, one in eight people, or approximately 853 million, would have diabetes, a 46% rise^[8]^.

Elevated triglycerides, decreased high-density lipoproteins, increased low-density lipoproteins, and elevated Apolipoprotein B are all components of a particular dyslipidaemia pattern ^[9]^. An assemblage of these irregularities in the same person appears to confer significant additional cardiovascular risk, beyond the risk associated with each aberration alone ^[8]^. Mild to moderate inflammation is a feature of Metabolic Syndrome, and C-reactive protein (CRP), the best-known measure of inflammation, can predict future cardiovascular events independently ^[10]^. MS patients with higher levels of high-sensitivity CRP have a higher risk of CVD, and CRP has been shown to interfere with insulin signalling and make atherothrombosis more likely ^[10]^. Studies have shown MS individuals to have raised amounts of CRP ^[11]^. Other studies, however, have shown that plant-based products can help lower CRP levels and other inflammatory biomarkers in people with Metabolic Syndrome ^[12]^. CRP measurement remains a good way to identify cardiac events in both healthy people and people who have coronary heart disease^[11]^. Many epidemiological and interventional studies have shown that good cardiorespiratory fitness (CRF) is linked to lower CRP levels, and that the link between CRF and cardiovascular events is mainly due to inflammatory factors ^[13]^. Furthermore, research has shown that oxidative stress is increased in chronic disease, where pro-inflammatory processes are not lacking.

Beta-caryophyllene (BCP) is the first naturally occurring dietary cannabinoid, a frequent component of the beneficial oil found in many spices and food plants, and an important constituent of cannabis. It exhibits a variety of anti-inflammatory properties in vivo, including cytoprotective, analgesic, and gastric anti-inflammatory effects ^[14, 15]^. The inflammation-reducing, insecticidal, and fungicidal qualities of BCP are well recognised ^[16]^. Plants including Piper nigrum, Cinnamon spp, and Origanum vulgare contain Beta-caryophyllene ^[17]^. The bark of Annona squamosa contains caryophyllene oxide, which demonstrates anti-inflammatory and analgesic properties ^[18]^. Beta-caryophyllene is a bicyclic sesquiterpene widely used in citrus flavours and spice blends, and as a preservative or additive (FDA-approved), intended to provide aroma to food and beverages. Due to its low toxicity, BCP has received the Generally Recognised as Safe (GRAS) designation ^[19]^.

The influence of diet on managing and improving chronic disease outcomes cannot be overemphasised. Several studies have explored the pharmacological activities of Beta-caryophyllene (BCP), including its strong anti-inflammatory properties, but no known study has explored the modulation of BCP on CVD indices in MS Wistar rats; hence, this study explored cardiovascular responses. Therefore, this study aimed to provide scientific evidence for the future therapeutic management of cardiovascular disease in MS using beta-caryophyllene. Beta-caryophyllene is relatively affordable when compared to many standard medications used for cardiovascular issues and disorders related to glucose and fat metabolism. Therefore, the impact on cardiovascular indices of dysfunction can be investigated as this study intended to achieve.

## Methods

Experimental procedures were performed in accordance with the “Guide for the Care and Use of Laboratory Animals” from the US National Institutes of Health ^[20]^. The experimental protocol was submitted and approved by the institution’s Animal Care and Use Research Ethics Committee (UI-ACUREC/17/0059) following international guidelines for animal research ^[20]^.

### Animals and experimental design

Seventy two (72) male Wistar rats (170-200 g), were maintained in a controlled temperature setting i.e room temperature (22± 1°C) with a controlled light–dark cycle (dark 06.00–18.00 hours), had unlimited access to water and food and given two weeks to get used to their new environment before the experiment commenced.

This study was performed in two major phases, with the rats being randomly allocated into 6 groups (n= 12) thus:

1. Group 1 (Normal Controls were administered a normal diet [ND])
2. Group 2 (ND+BCP).
3. Group 3 (HFHFD diet).
4. Group 4 (HFHFD continued +BCP).
5. Group 5 (HFHFD withdrawn). i.e returned to ND
6. Group 6 (HFHFD withdrawn +BCP). i.e returned to ND+ BCP.

Phase 1: Establishment of metabolic syndrome with a high-fat, high-fructose (HFHFD) diet administered for 16 weeks ^[5, 21]^, evidenced by hypertension, diabetes and obesity in the rats ^[22]^, via appropriate physiological measurements, i.e. anthropometry, mean fasting lipid and glucose profiles, and blood pressures; and Phase 2 was treatment with β-caryophyllene (BCP) (50 mg/kg, p.o., 4 weeks) ^[26, 27]^. Cardiovascular responses (C-reactive protein, apolipoprotein B, lipid profile, and oxidative stress) were measured in both research phases and in normal animals.

### Animal research diets

The standard diet (normal feed) comprised the following: Crude protein: 16.00%; Fats and oil: 5.0%; Crude fibre: 7.00%; Calcium: 1.60%; Available phosphorus: 0.45%; Lysine: 0.75%; Salt (min): 0.30%; Methionine: 0.36%; Kcal/kg metabolisable energy (min): 2450; net weight/kg: 25 g. The specific high-fat enriched diet administered was a modified mixture comprising 20% swine lard, 17% fructose, and 63% standard diet (normal feed)^[21]^. The high-fructose diet was administered via 10% fructose drinking water, freshly prepared daily using the weight/volume formula^[23]^.

### Source, dose selection, and mode of administration of Beta-caryophyllene (BCP)

BCP is a promising therapeutic option because of its broad availability, ease of use, and wide therapeutic window^[24]^. The (E)-β-caryophyllene used in this experiment was an 80% purified natural commercial sample (W225207) purchased from Sigma-Aldrich (St. Louis, Missouri, USA). Pure BCP, already in its liquid oily form, was suspended in corn oil as the vehicle for oral administration by gavage ^[25]^, at a dose of 50 mg/kg^[26, 27]^.

### Cardiac function assessments

Heart rate (HR), systolic and diastolic blood pressure (SBP and DBP), and mean arterial blood pressure (MABP) were measured in conscious, resting rats using a CODA Kent Scientific non-invasive blood pressure (NIBP) system. Rats were acclimatised to the restrainer before measurement. A tail cuff was applied, and the tail was maintained at 34–37°C using the system’s infrared heating platform to ensure adequate blood flow. We obtained ten measurements per rat and recorded the mean values. Electrocardiographic (ECG) measurements: Rats were anaesthetised intraperitoneally with ketamine/xylazine (KX; 0.1 mL/100 g body weight). After shaving and applying electrode gel, electrodes were placed on the limbs and chest and connected to an EDAN CARDIOMAQ PC ECG system. ECG parameters, including HR, P-wave duration, PR interval, QRS duration, QT interval, corrected QT (QTc; Bazett), and R-wave amplitude, were recorded^[28,29]^.

### Biochemical plasma analysis

Rats were fasted for six hours before blood collection. This time point was selected to prevent a postprandial glycaemic peak and an induced catabolic state in the rats^[30]^. Retro-orbital blood samples were collected in Microvette CB 300 Plasma/Lithium Heparin (Sarstedt, Germany) tubes. Samples were centrifuged for 10 minutes at 4°C at 3000 g [g = (1.118 x 10.5)R S2], where g is the relative centrifugal force (RCF), R is the rotor’s radius in centimetres, and S is the centrifuge’s speed in revolutions per minute^[30]^. Supernatants were collected, and, using the precise techniques outlined below, plasma concentrations of C-reactive protein, Apo-B, HDL, LDL, triglycerides, and cholesterol, as well as oxidative stress markers (total protein, reduced glutathione (GSH), superoxide dismutase (SOD), malondialdehyde (MDA), and nitric oxide), were assessed. As far as possible, all measurements were made from the same blood sample to avoid multiple fasting and blood collection periods^[30]^. The automated devices employed for the assays were as follows: 1) Molecular Devices Spectramax 190 Microplate Spectrophotometer; 2) Biobase ELI0A ELISA reader; 3) Centrifuge manufactured by Heraeus Instruments Biofuge Fresco; 4) Incubator manufactured by Awareness Technologies Stat Fax-2200.

### C - reactive protein, apolipoprotein-B, lipid profile and oxidative stress quantification

CRP: The Shanghai Bioassay Technology Laboratory’s RAT CRP/C-Reactive Protein/ELISA KIT measured high-sensitivity C-reactive protein. The Rat C-Reactive Protein (CRP) ELISA kit measures Rat CRP in blood, plasma, cell culture supernatants, and urine^[30, 31]^. Rat C-reactive protein was used as the immunogen; Total Cholesterol: this was analysed using a kit purchased from Fortress Diagnostics Limited (United Kingdom) ^[32]^, product code BXC0261; Triglycerides: Fortress Diagnostics Limited (United Kingdom)^[32]^ provided the Triglycerides kit, which was used to analyse this; the product code was BXC0271; HDL (High density lipoprotein): An HDL-cholesterol kit, product code BXC422A, from Fortress Diagnostics Limited (United Kingdom) was used for analysis; LDL (Low density lipoprotein) cholesterol: A homogeneous enzymatic selective protection technique for measuring serum and plasma levels of LDL cholesterol, i.e., an LDL-cholesterol kit (product code: BXC0431) from Fortress Diagnostics Limited (United Kingdom) was also used to analyse the LDL-cholesterol; Apolipoprotein B (Apo B): The Apolipoprotein B kit from Fortress Diagnostics, product code BXC0412, was used to perform this experiment. Assay procedures ^[33, 34, and 35]^ were used in tandem with the respective kit manuals.

Oxidative stress markers and their measurements included the following: 1) Determination of Protein using the Total Protein Kit (Fortress diagnostics, United Kingdom; Product code: BXC0173) (Biuret method)^[34, 35]^; 2) Reduced Glutathione (GSH) Assay Using the Glutathione Assay Kit. Reduced glutathione (GSH), which is made up of gamma-glutamyl, cysteine, and glycine, is the main free thiol tripeptide found in all live cells.; 3) Malondialdehyde (MDA) Assay Utilising the Lipid Peroxidation (MDA) Assay Kit ^[32, 34, 35^ ^]^: Lipid peroxidation refers to the deterioration of lipids due to oxidative damage and serves as an effective indicator of oxidative stress; 4) Superoxide Dismutase (SOD) Assay Utilizing the SOD Assay Kit ^[32, 34, 35^ ^]^: Superoxide dismutase (SOD) is a crucial antioxidative enzyme. Molecular oxygen and hydrogen peroxide are formed more easily when the superoxide anion (O2.-) breaks down with the aid of this enzyme; 5) Nitric Oxide Assay utilising the Non-Enzymatic Colorimetric Method for Nitric Oxide: this was established using the assay kit for non-enzymatic nitric oxide (Product Number: NB88) produced by Oxford Biomedical Research, Inc., Oxford, MI 48371, U.S.A^[36]^. The nitric oxide kit measured the total amount of nitric oxide (NO) produced in lab experiments after metallic cadmium converted nitrate to nitrite. Spectrophotometry may be utilised to evaluate the amount of nitric oxide by measuring the quantity of its steady breakdown products, nitrate and nitrite. The two substances, nitrate (NO3-) and nitrite (NO2-), are produced when nitric oxide (NO) is oxidised. They have been measured in blood to show how active the nitric oxide synthase enzyme of endothelium is, as well as to get a rough idea of NO levels. The Griess reaction^[37]^ is then used to measure nitrite. The kit can measure the total amount of nitric oxide in samples with high protein content. Compared to nitrate reductase, the cadmium catalyst is much more stable in the presence of harsh chemicals that break down proteins. This kit can accurately measure nitric oxide levels as low as 1 pmol/mL (∼1 µM) in water. A minimum sample amount of one hundred microliters is needed, and it depends on the quantity of NO in the sample. 540 nm was used to measure the finished process ^[(36, 38]^.

### End of experiment

This consisted of animal euthanasia and sacrifice, harvesting and weighing of animal organs for histopathological analyses of the tissues.

### Statistical analyses

The Statistical Package for the Social Sciences (SPSS) version 23 was used for data analyses. Data summarisation was performed using means and standard deviations; bar charts were plotted using GraphPad Prism 18. A paired t-test compared the groups under investigation. ANOVA (Analysis of Variance) was used to assess post-treatment parameters when there were more than two groups. An independent t-test was used to compare two groups. A post hoc pairwise comparison of group means was performed after ANOVA. The significance level for each test was set at 5%, and P values and 95% confidence intervals were provided for significance tests.

## Results

### Cardiac-electrophysiology

Table 1 showed that systolic blood pressure, rate, flow, and volume in the HF/HFD-Withdrawn group decreased by 20%, 35%, 85%, and 20%, respectively, compared with the HF/HFD-Continued group. The heart rate increased significantly by 51% in the HF/HFD-Withdrawn+BCP relative to the HF/HFD-Continued+BCP, whereas volume decreased significantly by 39% in the HF/HFD-Continued+BCP relative to the HF/HFD-Withdrawn+BCP.

**Table 1:** Modulatory Outcome of BCP treatment on Heart rate and Blood Pressure (BP) Variables among HFHFD animal sets on HFHD diet).

|  | HF/HFD Continued | HF/HFD-Continued+BCP | HF/HFD-Withdrawn | HF/HFD-Withdrawn+BCP |
| --- | --- | --- | --- | --- |
| Systolic (mmHg) | 145.0±12.13 | 120.3±2.15 | 129.0±1.28* | 125.5±11.1 |
| Diastolic (mmHg) | 105.7±10.8 | 94.0±17.2 | 91.50±9.35* | 95.5±11.2 |
| MABP (mmHg) | 114±7.09 | 117±10.3 | 116±11.38 | 104±3.14 |
| Heart Rate (/min) | 448±11.9 | 280±18.7 | 380±16.2* | 351±3.00** |
| Flow rate (ml/min) | 19.60±3.91 | 14.71±11.38 | 24.3±0.47 | 13.93±6.20 |
| Volume (ml/kg) | 69.2±5.10 | 68.0±4.40 | 82.5±1.50 | 49.3±6.41** |

Data was displayed by means with a standard deviation of 12 animals. \*= substantially distinct from HFHFD-continued, \*\*= substantially distinct from HFHFD-withdrawn. HF/HFD= High fat and High fructose diet, BCP= Beta Caryophyllene

Table 2 showed that HR and PR interval increased significantly in the HF/HFD-Continued+BCP group compared with the HF/HFD-Continued group. Specifically, HR and PR interval increased by 35% and 27%, respectively. In the HF/HFD-Withdrawn+BCP group, HR and PR interval decreased. In addition, the QT segment, QTc and R amplitude increased significantly by 22%, 31% and 58%, respectively, in the HF/HFD-Withdrawn+BCP group compared with the HF/HFD-Withdrawn group.

**Table 2:** Modulatory Outcome of Beta Caryophyllene (BCP) on ECG Variables among HFIIFD-Rats on IIFIIFD.

|  | HF/HFD Continued | HF/HFD-Continued+BCP | HF/HFD-Withdrawn | HF/HFD-Withdrawn+BCP |
| --- | --- | --- | --- | --- |
| HR (/min) | 190.0±55.0 | 140.0±47.0* | 258.0±5.45 | 252.5±23.0 |
| P duration (Ms) | 24.0±1.15 | 25.0±6.20 | 19.5±2.12 | 24.50±0.50 |
| PR interval (Ms) | 43.0±3.50 | 55.0±15.60* | 49.5±3.53 | 45.0±12.70 |
| QRS duration (Ms) | 13.0±1.4 | 14.00±6.0 | 14.0±0.56 | 17.0±1.40 |
| QT segment (Ms) | 83.0±39.6 | 76.7±23.0 | 70.5±6.36 | 86.0±18.4** |
| QTc (Bazett) | 115.5±19.0 | 111.7±13.0 | 145.5±12.0 | 176.5±6.4** |
| R amplitude (mV) | 0.45±0.025 | 0.30±0.005 | 0.367±0.01 | 0.58±0.024** |

Data was displayed by means with standard deviation of 12 animals. \*= substantially distinct from HFHFD-continued,
\*\*= substantially distinct from HFHFD-withdrawn. HF/HFD= High fat and High fructose diet, BCP= Beta Caryophyllene.

### Electrocardiography (ECG)

Animals regularly fed a high-fat, high-sugar diet showed clear changes in cardiac activity, as seen in ECG tests. The tests showed problems with the heart’s rhythm, including abnormal beating in both the upper and lower chambers. They also indicated prolonged PR intervals, which suggests heart damage.

Animals whose meals were changed (i.e., returned to a normal diet) and given Beta-caryophyllene showed better results in the traits observed on the ECG.

Animals that continued to eat a high-fat, high-fructose diet showed different ECG results compared with those whose meals were stopped during Beta-caryophyllene treatment.

### C-reactive protein

Fig 1 shows an increase in CRP levels on the Metabolic Syndrome diet, which was ameliorated by BCP. BCP, however, decreased CRP levels, i.e., between HF/HFD-continued+BCP and HF/HFD-continued, as well as between HF/HFD-withdrawn+BCP and HF/HFD-withdrawn. However, these differences were not statistically significant.

**Fig 1:**
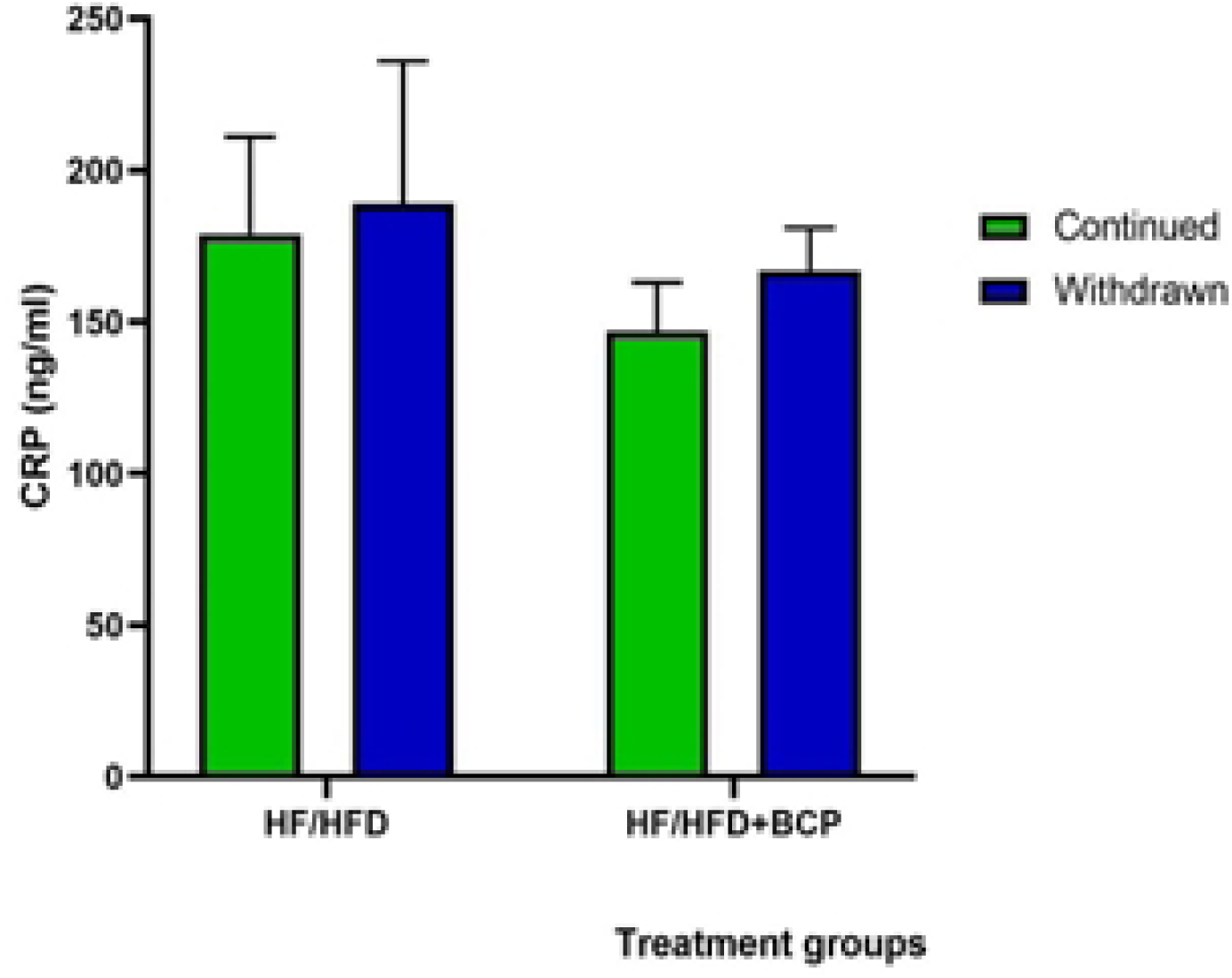
Modulatory Outcome of BCP on CRP Level of Animals Administered HFHFD. Values were displayed *as means and standard deviation of 12 animals. HF/HFD= High fat and High-fructose diet. BCP = Beta-caryophyllene*.

### Apolipoprotein-B

In Fig 2, APO B levels increased substantially (p<0.05) across animal groups exposed to both the HF/HFD-continued and withdrawn diets. However, APO B levels decreased in the BCP treatment groups, i.e., HF/HFD withdrawn + BCP, compared with HF/HFD continued + BCP.

**Fig 2:**
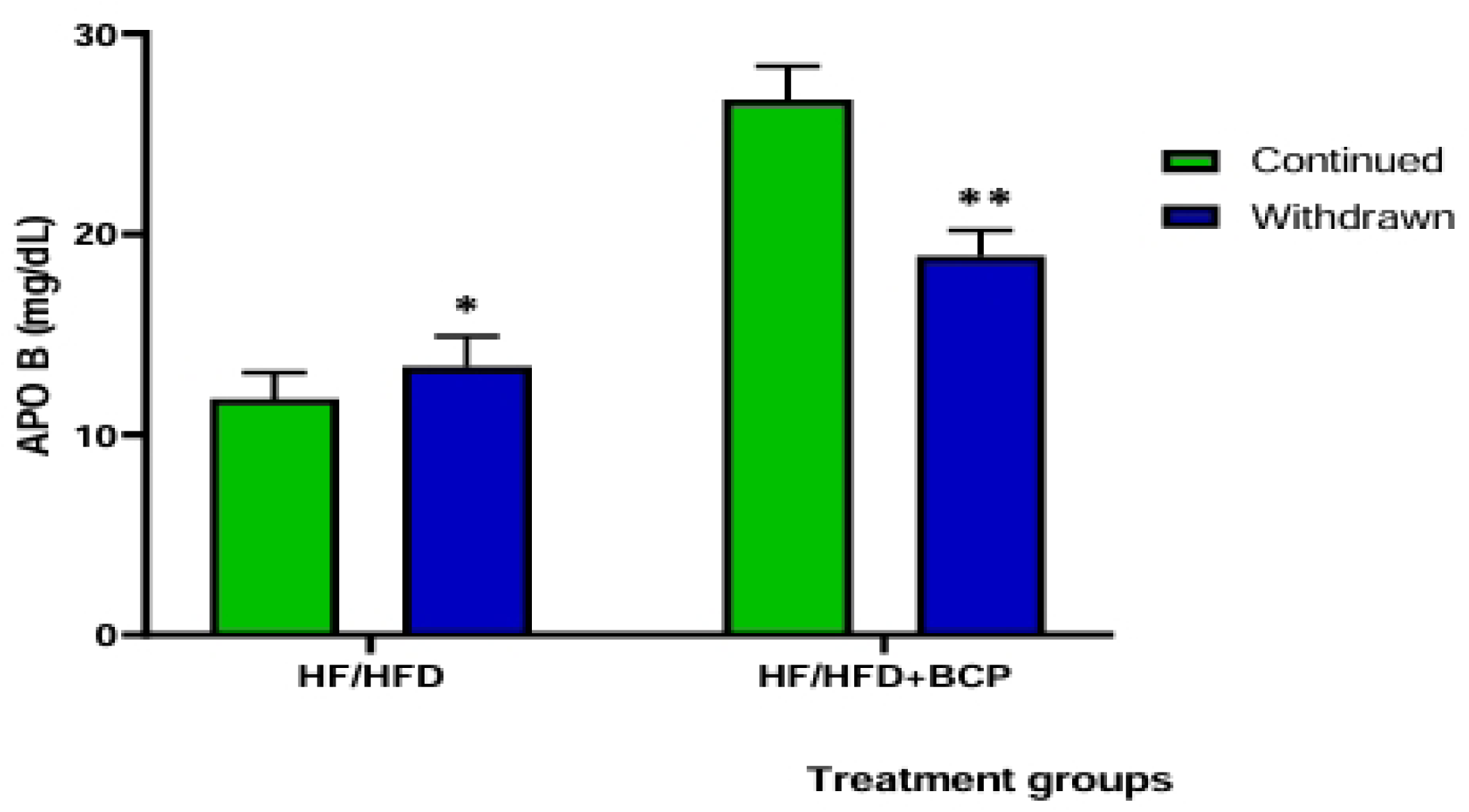
Modulatory Outcome of BCP on Serum Apolipoprotein B (APO B) Level of Rats Fed on HFHFD. *Values were presented as means and standard deviations For 12 animals. *-substantially different from HF/HFD-continued; **= substantially different from HF/HFD-withdrawn. HF/HFD = high-fat, high-fructose diet; BCP = beta-caryophyllene*

### Lipid profile

In Fig 3, animal groups exposed to HFHFD-continued+BCP had significantly lower levels of total cholesterol (TC) than those subjected to HFHFD-continued. Additionally, HFHFD-withdrawn+BCP significantly decreased TC (p<0.05) compared with HFHFD-withdrawn.

**Fig 3:**
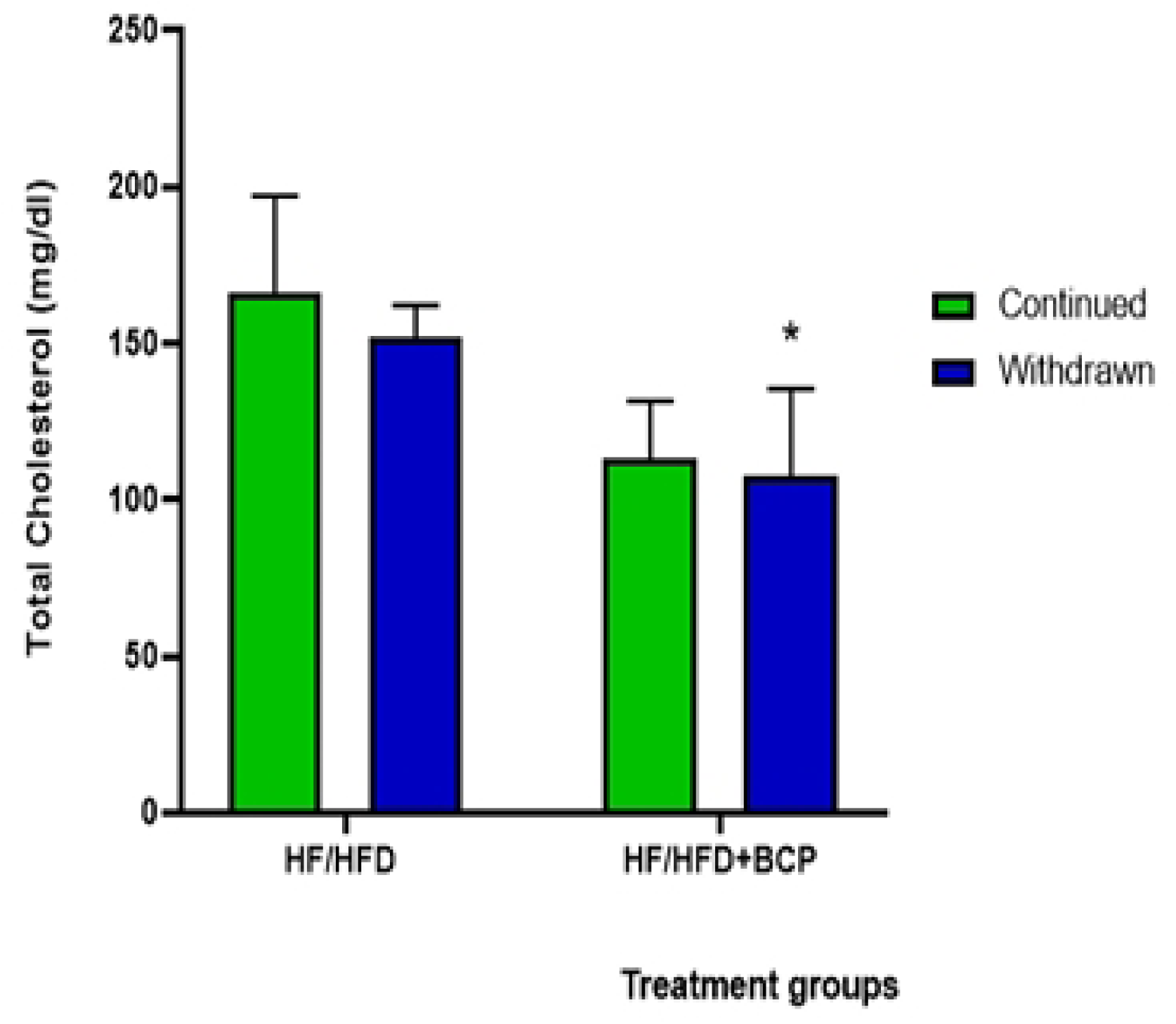
Modulatory Outcome of BCP on serum Total Cholesterol.

In Fig 4, administration of Beta Caryophyllene led to a substantial rise in HDL (p<0.05) among animals that had their HFHFD diets withdrawn, i.e., HFHFD-withdrawn+BCP, compared with withdrawal of the HFHFD without BCP.

**Fig 4:**
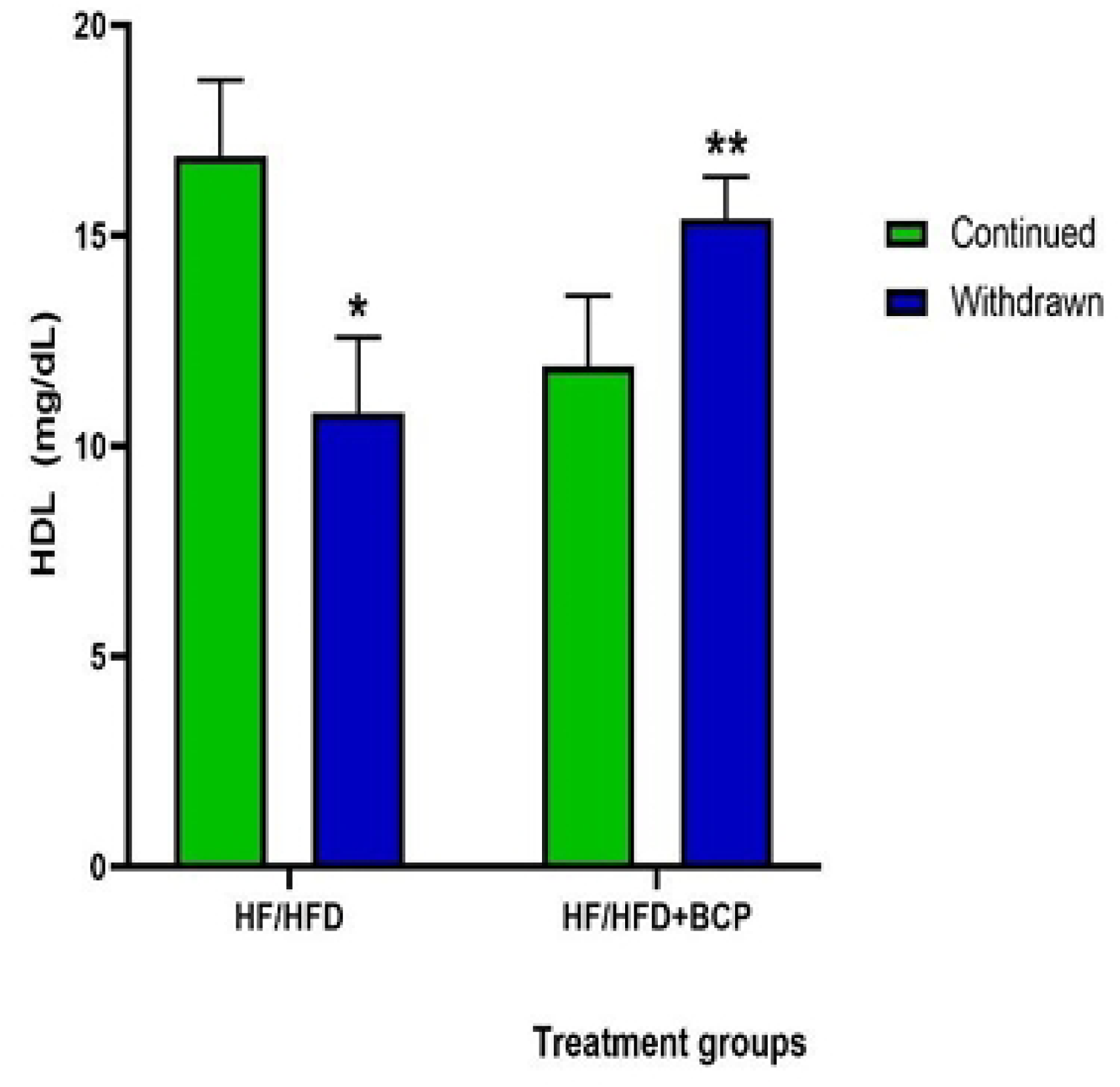
Modulatory Outcome of BCP on Serum High Density Lipoprotein (HDL) of Rats Fed on HFHFD.

Fig 5 shows decreased levels of low-density lipoprotein (LDL) among animals exposed to HFHFD withdrawn + BCP, compared with those on HFHFD continued + BCP. However, without BCP administration, LDL increased significantly (p<0.05) among animals that had their HFHFD

**Fig 5:**
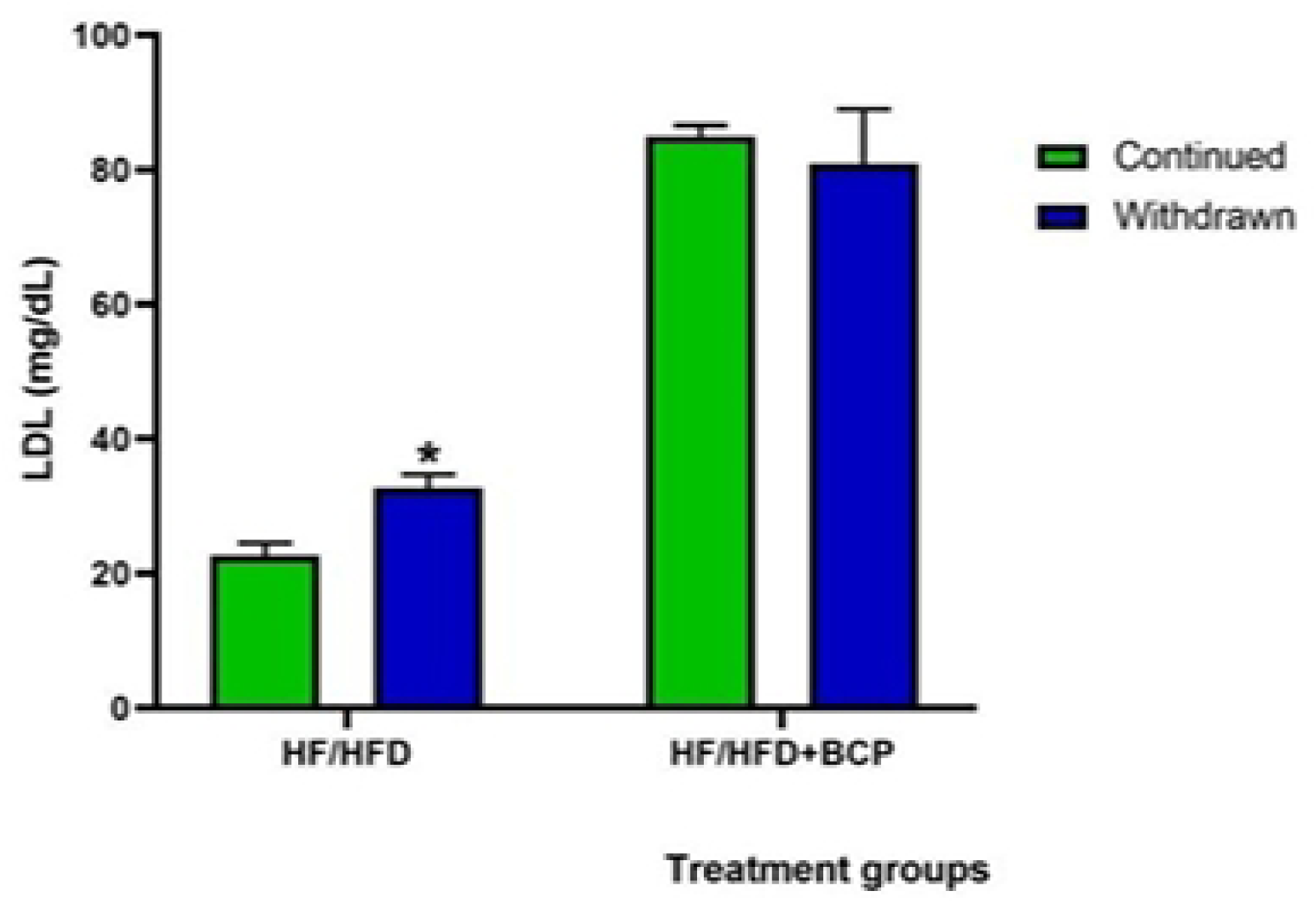
Modulatory outcome of β-caryophyllene (DCF) on serum low density lipoprotein (LDL) of rats fed on HFHFD.

Fig 6 shows that TG levels decreased significantly (p<0.05) among animals whose HFHFD was withdrawn, compared with those continuing on HFHFD. BCP was observed to reduce TG levels among animals on withdrawn HFHFD+BCP compared with those continuing on HFHFD+BCP.

**Fig 6:**
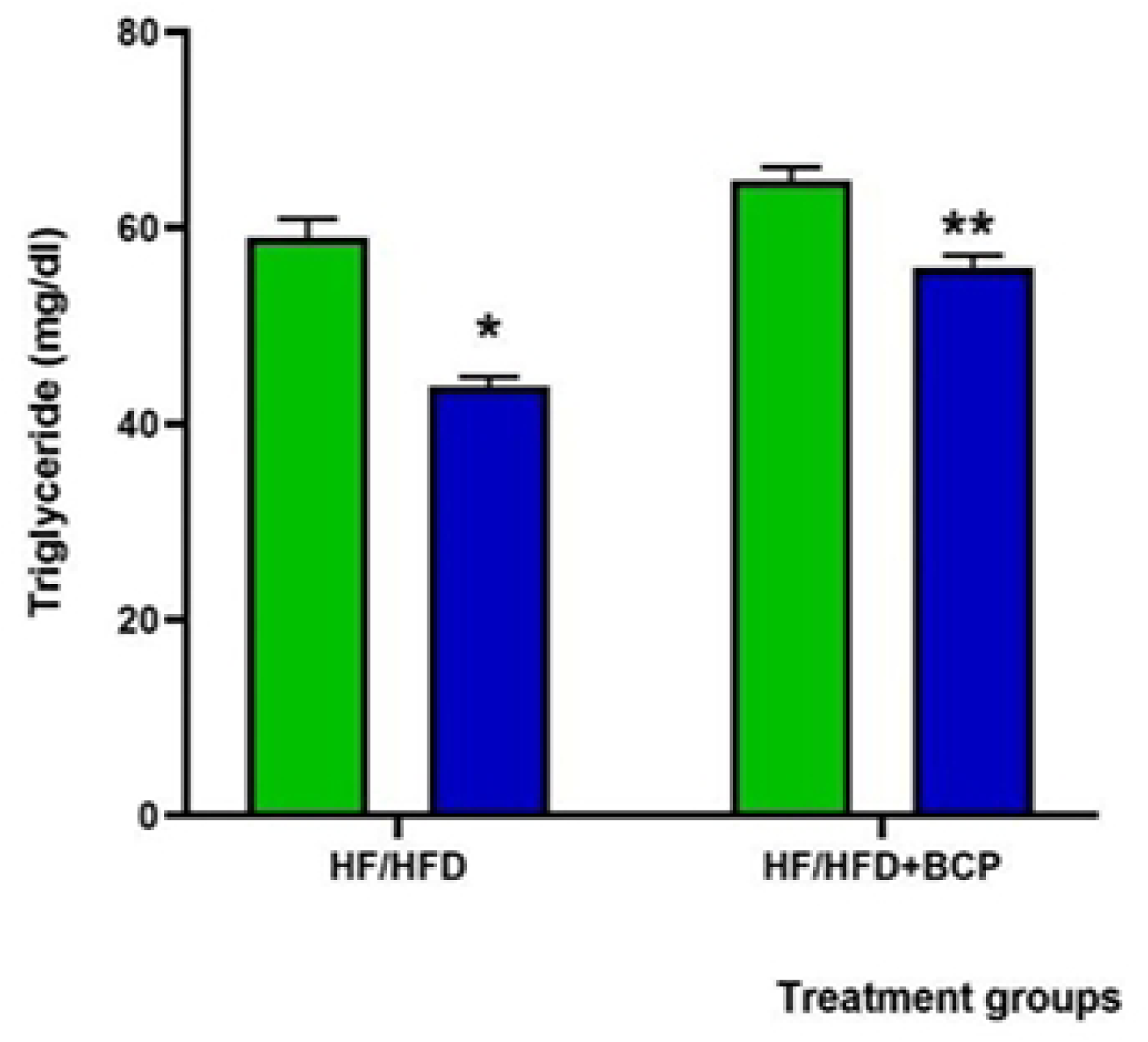
Modulatory Outcome of BCP on Serum Triglyceride Level of Rats Fed on HFHFD.

### Oxidative stress

Fig 7 shows a decrease in protein levels in the HF/HFD-continued+BCP group compared with HF/HFD-continued, as well as in the HF/HFD-withdrawn+BCP group compared with HF/HFD-withdrawn. Specifically, the HF/HFD-continued+BCP and HF/HFD-withdrawn+BCP groups decreased serum protein levels by 24% and 34%, respectively.

**Fig 7:**
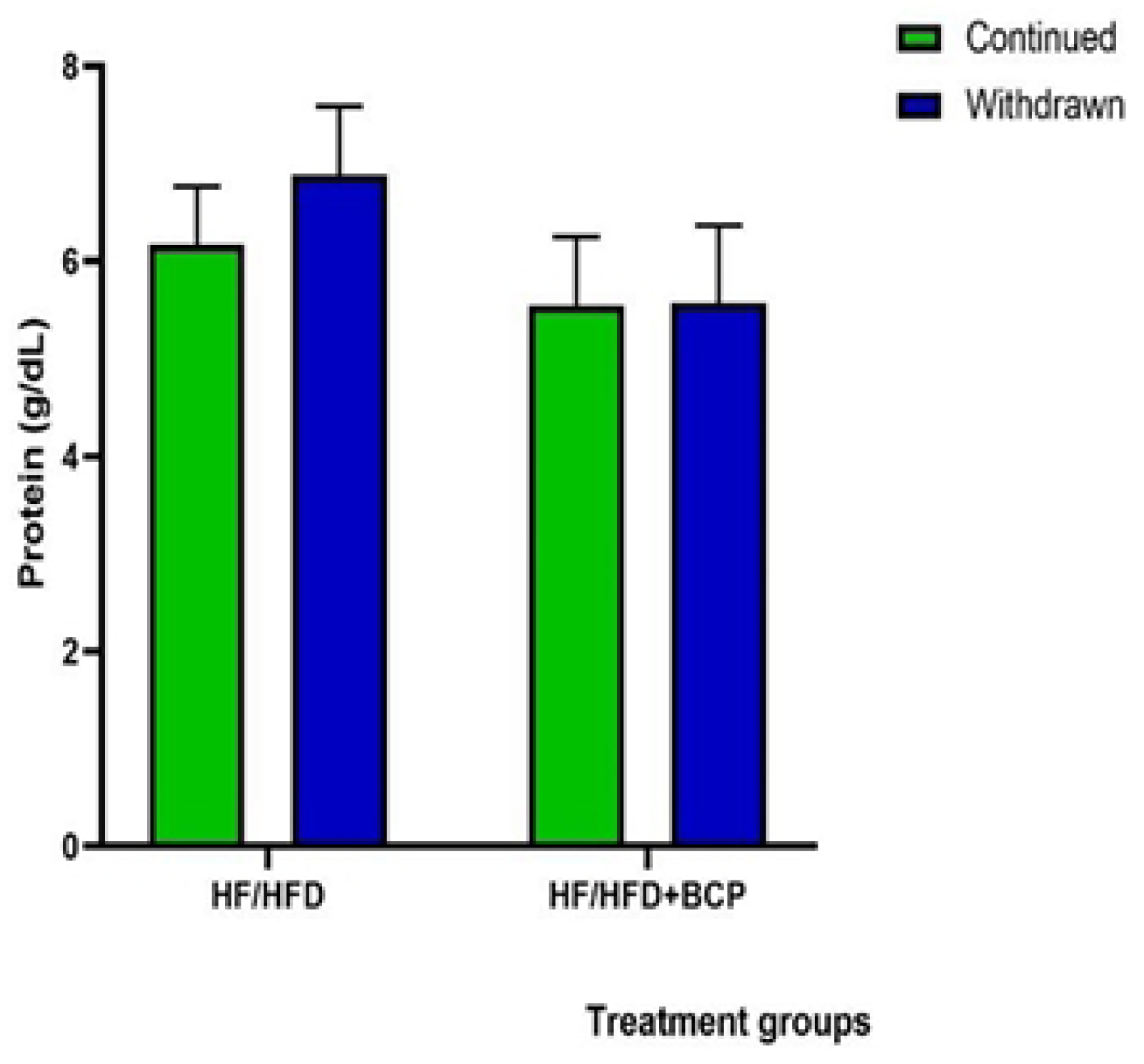
Modulatory outcome of BCP on mean serum protein levels among animals administered HFHFD.

Fig 8 shows that reduced glutathione levels increased significantly in the HF/HFD-continued+BCP group compared with the HF/HFD-continued-only group. In this group, GSH levels increased by 44%. Likewise, GSH levels increased significantly in the HF/HFD-withdrawn+BCP group compared with the HF/HFD-withdrawn group. BCP appreciably increased GSH levels in the HF/HFD-withdrawn diet+BCP group.

**Fig 8:**
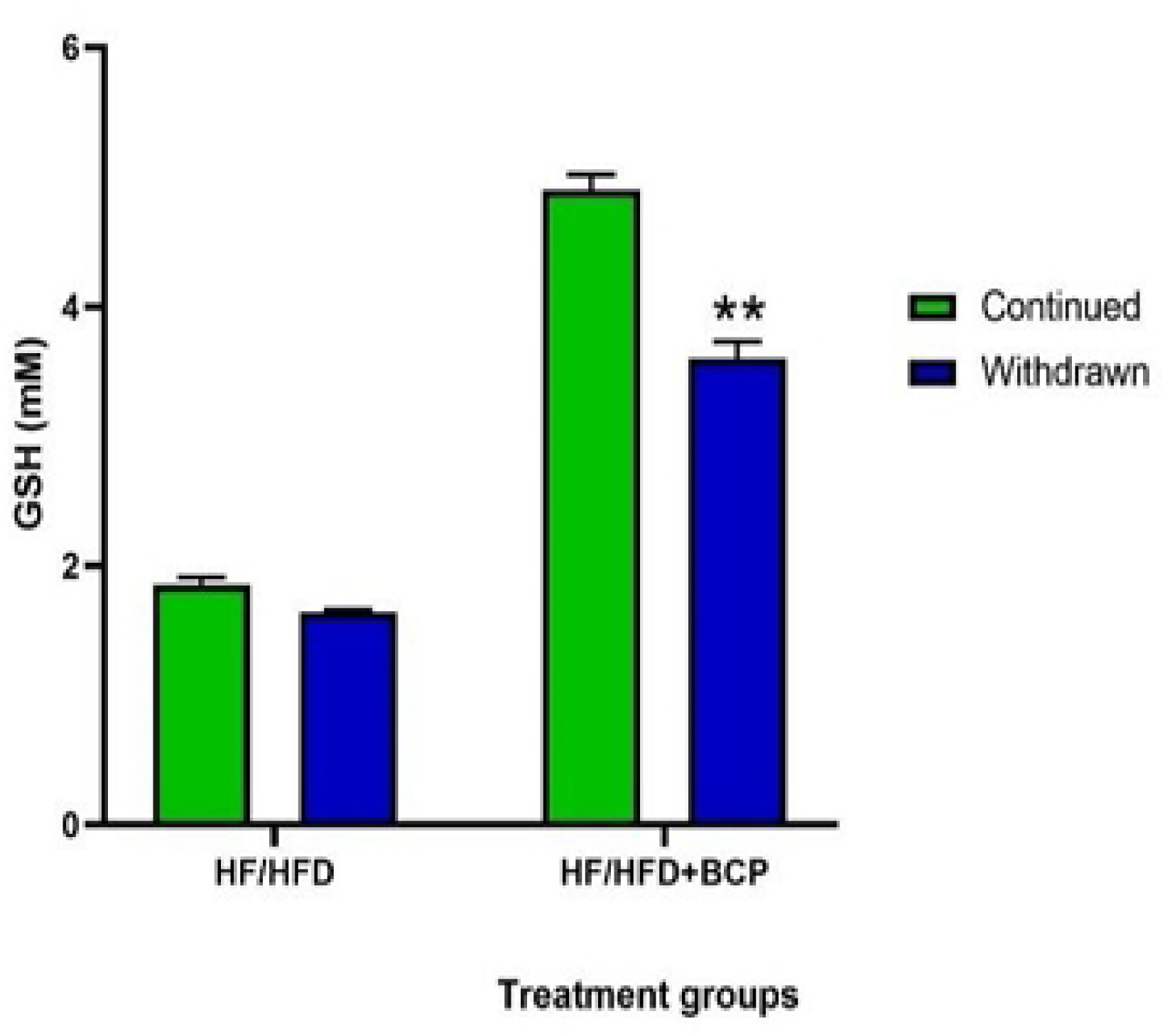
Modulatory outcome of BCP upon mean GSH levels among animals administered HFHFD.

The results in Fig 9 indicate that superoxide dismutase activity increased substantially in the HF/HFD-continued+BCP group (p<0.05) compared with HF/HFD-continued, and in the HF/HFD-withdrawn+BCP group compared with HF/HFD-withdrawn. Specifically, Beta Caryophyllene increased SOD activity by 67% in HF/HFD-continued+BCP and by 64% in HF/HFD-withdrawn+BCP.

**Fig 9:**
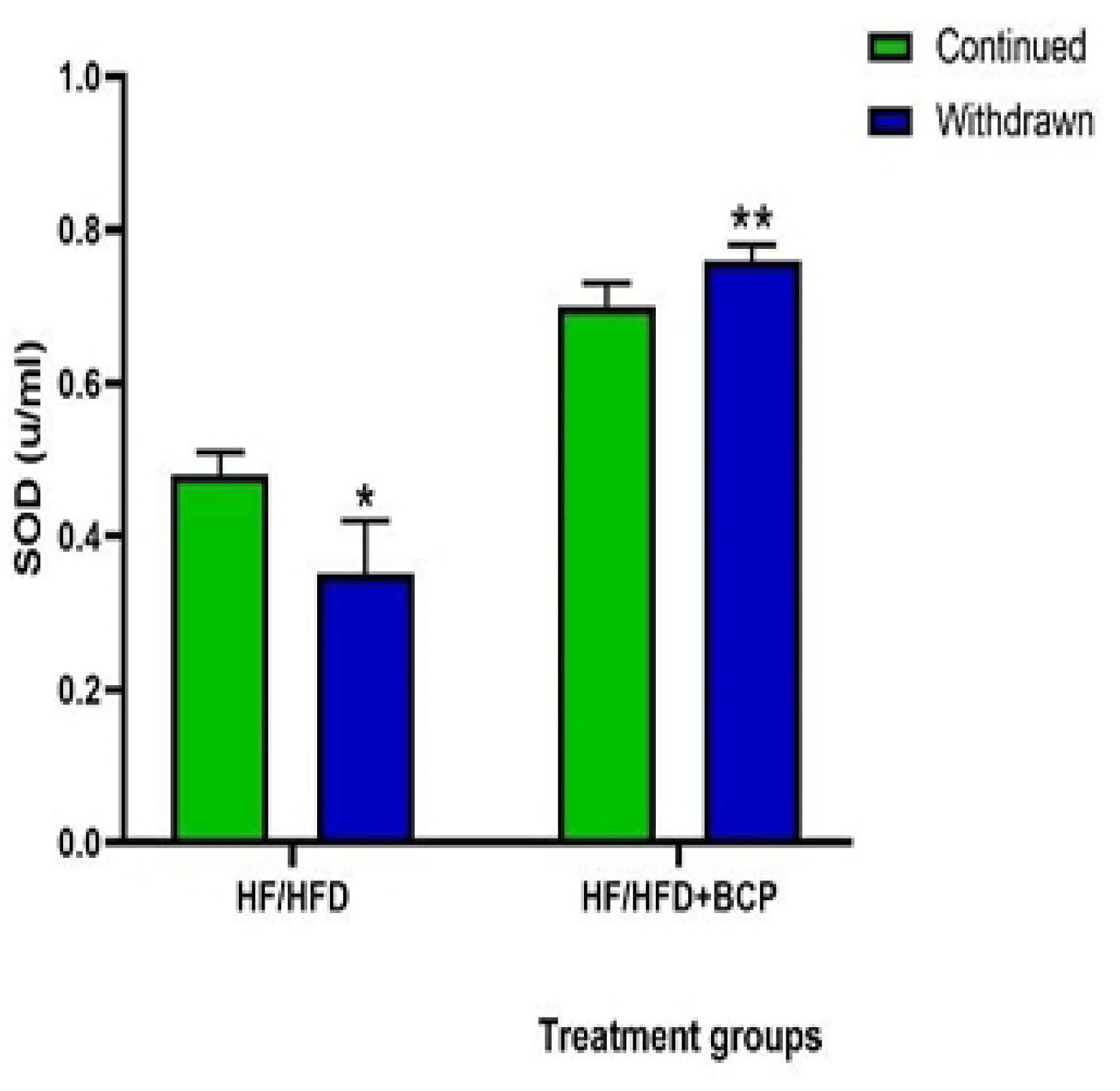
Modulatory outcome of BCP on Superoxide Dismutase (SOD) activity among animals on HFHFD.

In Fig 10, animals exposed to HF/HFD-withdrawn+BCP had markedly lower malondialdehyde levels (p **≤** 0.05) than those subjected to HF/HFD-continued+BCP. MDA decreased by 19%.

**Fig 10:**
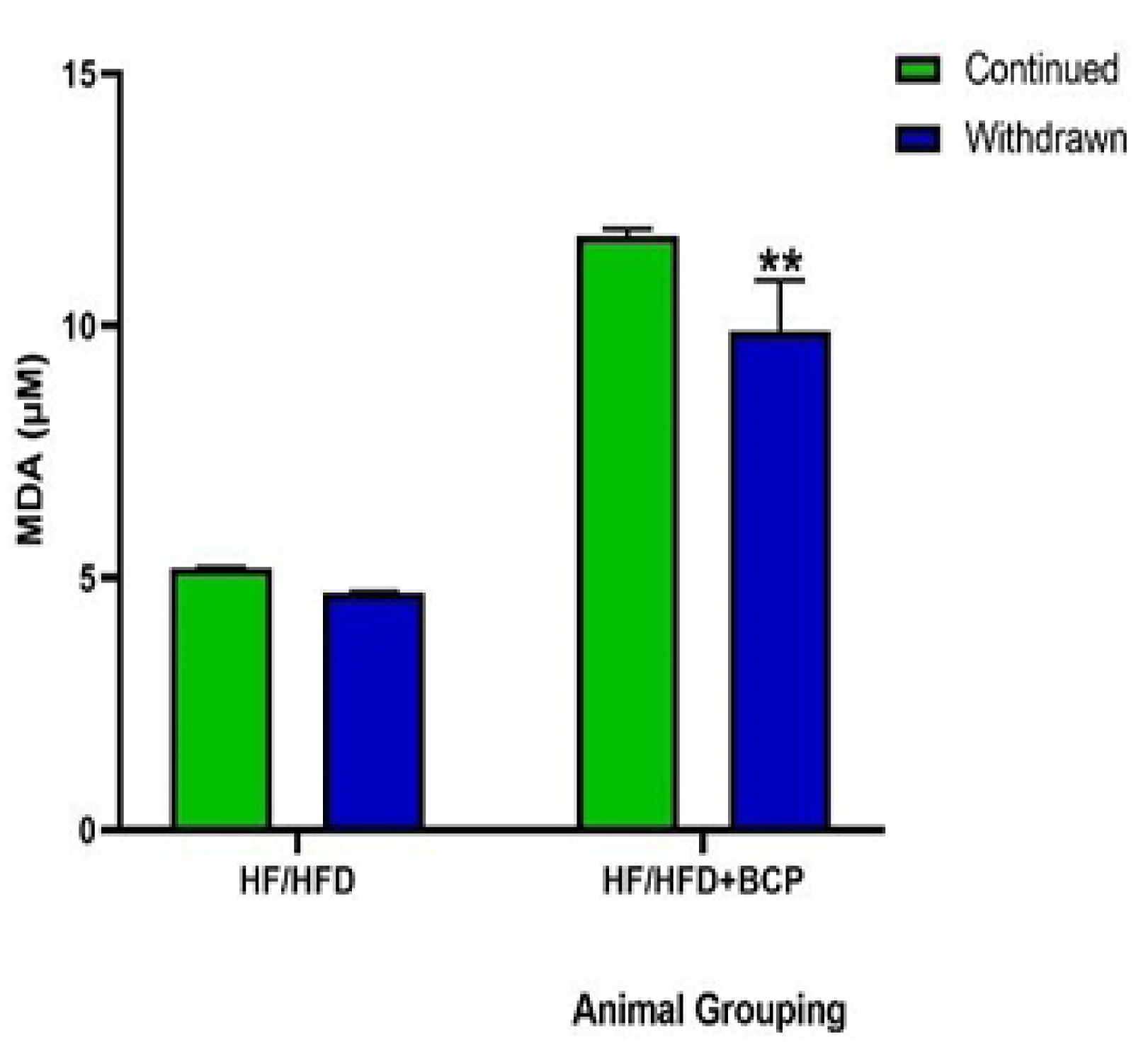
Modulatory outcome of BCP on Malondialdehyde (MDA) levels among animals administered HFHFD.

Figure 11 shows a substantial increase in nitric oxide levels (p<0.05) in the HF/HFD-continued+BCP group compared with the HF/HFD-continued group, as well as in the HF/HFD-withdrawn+BCP group compared with the HF/HFD-withdrawn group. However, nitric oxide levels were lower in the HF/HFD-withdrawn+BCP group than in the HF/HFD-continued+BCP group.

**Fig 11:**
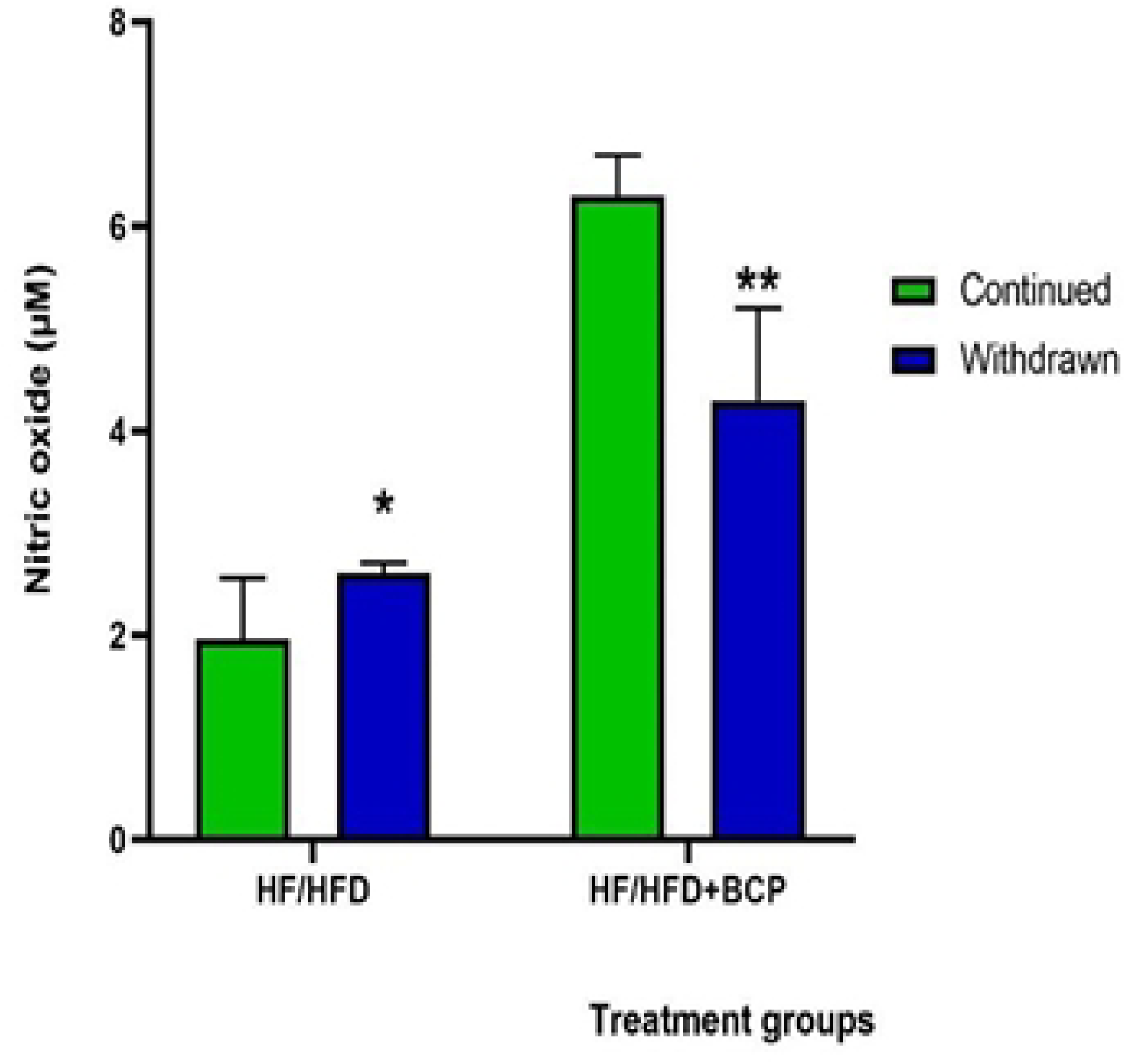
Modulatory outcome BCP on Nitric oxide level of animals administered HFHFD.

**Photomicrographs (in S12-14 figs section)**

## Discussion

### Cardiovascular and haemodynamic parameters

The World Health Organisation identifies raised blood pressure (hypertension) as a major contributor to cardiovascular mortality, accounting for approximately 16.5% of deaths globally each year^[39]^. In the present study, β-caryophyllene (BCP) improved several cardiovascular parameters in rats with metabolic syndrome. Treatment reduced systolic blood pressure (SBP) from 145.00±12.13 to 120.30±2.15 mmHg, diastolic blood pressure (DBP) from 105.70±10.80 to 94.00±17.20 mmHg, and heart rate (HR) from 448.00±11.90 to 280.00±18.70 beats/min. These findings are relevant, as hypertension is a major risk factor for cardiovascular morbidity and mortality^[39]^. Proposed mechanisms underlying hypertension include vascular smooth muscle proliferation and vasoconstriction, increased intracellular calcium, enhanced arterial reactivity, oxidative stress, and impaired endothelial nitric oxide synthase activity^[40]^. Activation of the renin–angiotensin–aldosterone system may further contribute to elevated blood pressure^[41]^.

Withdrawal of the HF/HFD regimen was associated with a 20% reduction in SBP, a 35% reduction in HR, an 85% reduction in blood flow, and a 20% reduction in blood volume compared with continued HF/HFD exposure. Among BCP-treated animals, blood volume was 39% lower in the HF/HFD-continued+BCP group than in the HF/HFD-withdrawn+BCP group, while HR was 51% higher following diet withdrawal. These findings suggest that both dietary modification and BCP may contribute to improved cardiovascular function. Previous evidence supports a vasorelaxant effect of Annona squamosa seed constituents through inhibition of extracellular calcium influx via voltage-dependent calcium channels ^[42,43,44]^. Similarly, BCP-rich Eugenia punicifolia preparations have been reported to reduce systolic and diastolic blood pressure and improve other metabolic and inflammatory parameters ^[45.]^

The improvement in DBP observed after HF/HFD withdrawal and BCP treatment is consistent with evidence that dietary modification can reduce blood pressure ^[46]^. Other plant-derived preparations have also demonstrated antihypertensive effects; for example, Melothria maderaspatana tea reportedly reduced SBP and DBP by 23.8 and 15.5 mmHg, respectively, after 45 days in mildly hypertensive individuals ^[47]^.

BCP also reduced mean arterial blood pressure (MABP) in the present study. This finding aligns with previous research showing significant reductions in SBP, DBP and MABP after administration of Persea americana and Annona muricata leaf preparations containing BCP^[^**⁴**²^]^. The authors proposed inhibition of the renin–angiotensin–aldosterone pathway as a possible mechanism.^[^**⁴⁸**^]^ The improvement in HR following HF/HFD withdrawal and BCP treatment may also reflect improved endothelial and cardiovascular function. Similar effects have been reported for Bursera simaruba, which reduced HR and blood pressure while enhancing endothelial nitric oxide synthase activity ^[^**⁴⁹**^]^

Changes in blood flow and volume further suggest improved cardiovascular function after BCP treatment. Abnormal blood flow and altered endothelial shear stress can promote endothelial dysfunction and vascular injury ^[50,51]^. In the present study, blood flow and volume increased after withdrawal of the HF/HFD diet in BCP-treated animals, suggesting possible improvements in cardiac output and peripheral vascular function. Comparable cardiovascular effects have been reported with other medicinal plant preparations ^[48,49]^.

### Electrocardiographic findings

ECG assessment demonstrated significant electrophysiological abnormalities in rats exposed to the HF/HFD regimen, including alterations in HR, P-wave duration, PR interval and QT interval. These findings suggest impaired cardiac electrical function associated with metabolic syndrome. Importantly, BCP treatment modified several of these abnormalities. HR and PR interval increased significantly in the HF/HFD-continued+BCP group compared with the untreated HF/HFD-continued group, while QT interval, QTc and R-wave amplitude increased by 22%, 31% and 58%, respectively, in the HF/HFD-withdrawn+BCP group compared with the HF/HFD-withdrawn group.

The prolonged ECG intervals observed after HF/HFD exposure may indicate underlying myocardial injury or altered cardiac conduction. The QT interval spans ventricular depolarisation and repolarisation and is influenced by the coordinated movement of sodium, calcium and potassium ions across myocardial cell membranes^[52]^. Dysfunction of these ion channels can prolong ventricular repolarisation and the QT interval^[52]^. In the present study, BCP reduced P-wave duration in animals after diet withdrawal, although this difference was not statistically significant. Overall, the findings suggest a modulatory effect of BCP on cardiac electrophysiology, with no evidence of clinically concerning QT prolongation.

The present findings are consistent with previous experimental evidence that metabolic disorders can produce early cardiac electrical abnormalities before overt structural changes. In diabetic rats fed a high-fat diet and treated with streptozotocin, alterations in cardiac electrical conduction preceded structural changes, including hypertrophy^[53]^. Increased T-wave amplitude, ST-segment height and QRS duration have also been associated with ventricular repolarisation abnormalities and increased cardiovascular risk in such models^[53,54]^. These observations support the possibility that the ECG abnormalities observed in the present study reflect early myocardial and conduction-system effects of metabolic syndrome.

Atrial fibrillation was also observed in rats with metabolic syndrome but was reversed after BCP treatment. Atrial fibrillation may arise from structural and electrical remodelling, including atrial enlargement and impaired conduction^[55]^. The reversal observed with BCP treatment further suggests a potential cardioprotective and electrophysiological modulatory effect of BCP, thereby contributing to improved overall cardiorespiratory function.

### C-reactive protein (CRP)

Individuals with Metabolic syndrome already have elevated inflammation and CRP levels. Higher levels of high-sensitivity CRP are associated with a greater risk of cardiovascular disease. CRP has been shown to interfere with insulin signalling, which makes atherothrombosis more likely^[10]^, ^[11]^. Other studies, however, have shown that plant-based products can help lower CRP levels and other inflammatory biomarkers in people with Metabolic syndrome^[12]^. CRP measurement remains a good way to identify cardiac events in both healthy people and those with coronary heart disease^[13]^.

Recently, many epidemiological and interventional studies have shown that good cardiorespiratory fitness (CRF) is linked to lower CRP levels. The link between CRF and cardiovascular events is largely mediated by inflammatory factors, and research has demonstrated that overweight individuals with MS had higher levels of CRP^[13]^. These higher levels were significantly correlated with measures of sympathetic and parasympathetic activity compared with control groups^[13]^.

The current study investigated how β-caryophyllene affected cardiac dysfunction in Wistar rats with Metabolic Syndrome induced by the HFHFD. The HFHFD caused Wistar rats to develop Metabolic Syndrome, which raised CRP levels in both the groups that continued eating and the ones that stopped the diet. When BCP was given, however, there was a clear drop in CRP levels in the HF/HFD continued+BCP groups compared with HF/HFD-continued, and in the HFHFD-withdrawn+BCP groups compared with HF/HFD withdrawn. These differences, however, were not statistically significant.

Similarly, another study examined serum levels of inflammatory markers in rats that had been administered alloxan and then treated with aqueous leaf extract of Terminalia catappa and insulin from outside the body. It reported a substantial rise in serum CRP, Interleukin-6, and fibrinogen in the diabetic group, in contrast to the control group. In the diabetic group given the extract and insulin, inflammatory biomarkers decreased substantially. The team concluded that aqueous leaf extract of Terminalia catappa could lower some inflammatory markers and improve inflammation in diabetes in a way that was similar to insulin from outside the body^[56]^. As previously stated, β-Caryophyllene occurs in significant amounts in some medicinal plants.

In an original study^[57]^, intraperitoneal administration of the ethanolic extract of Nigella sativa seeds significantly reduced blood CRP levels (p<0.001) compared with the control group. The ethanolic extract of Nigella sativa significantly lowered CRP levels (p<0.05), more than diclofenac sodium^[57]^.

### Lipid profile and Apolipoprotein –B (Apo-B)

Current research indicates that individuals with low-to-moderate dyslipidaemia who have low cardiovascular risk may benefit from using certain clinically active nutraceuticals that have been shown to lower lipids. These include plant sterols and stanols, soluble fibres, and green tea extracts, among others, to safely improve their plasma lipid levels^[58]^.

Research evidence^[59]^ has documented more than seventy therapeutic plants shown to lower blood lipid levels and offer marked advantages in effectiveness, safety, affordability, and acceptability^[59]^. As noted earlier, many medicinal plants contain β-caryophyllene. The use of plants for therapy also occurs in urban areas in developed countries^[59]^. Furthermore, a vast body of recent research from Africa and other parts of the globe has shown that certain plant extracts are beneficial in ameliorating dyslipidaemia, thereby reducing the development of CVD. For example, a study conducted in Ethiopia^[60]^ found that M. africana leaf extract (methanolic extract of Myrsine africana) possesses antidyslipidaemic as well as antidiabetic properties. This was demonstrated by higher HDL cholesterol, hexokinase, and insulin, as well as reductions in blood glucose, glycated haemoglobin, glucose-6-phosphatase, fructose-1,6-bisphosphatase, total cholesterol, and triglycerides^[60]^.

#### Total cholesterol (TC)

The risk of cardiovascular death is significantly increased by lipid disorders^[61]^. Metabolic Syndrome predicts the development of CVD, and behavioural risk factors such as poor diet, inactivity, tobacco use, and excessive alcohol consumption are linked to over 80% of CVD cases^[61]^. The consequences of this poor diet may manifest as obesity, overweight, and elevated blood lipids. Moreover, lipid disorders are the most common modifiable risk factors^[61]^.

According to a Polish study assessing lipid profiles in individuals with Metabolic Syndrome, cardiovascular risk was evaluated using the SCORE chart^[61]^. The study found that patients with MS had an extremely significant cardiovascular risk. Compared with the high- and medium-CV-risk groups, the MS group had markedly higher levels of LDL-C, TG, and non-HDL-C44. The study concluded that elevated TC levels and other lipid fractions may be linked to elevated CV risk in this patient population^[61]^.

Individuals diagnosed with Metabolic Syndrome (Ms) have a heightened risk of developing CVD and display several lipid abnormalities, including raised LDL-c ^[62]^. These include reduced HDL-c, increased triglycerides, and small, dense LDL (sd-LDL) particles ^[62]^. Nonetheless, a different study in men found that every component of the lipid profile was linked to multiple sclerosis, and that preventing dyslipidaemia and aiming for a balanced lipid profile would likely be effective measures to lower its incidence ^[63]^.

The current research results revealed that BCP substantially decreased total cholesterol levels in Wistar rats in the withdrawn and continuing HFHFD groups compared with the corresponding controls. This demonstrates that BCP treatment improved total cholesterol, with the effect more pronounced after HFHFD withdrawal.

Similarly, various plant extracts high in flavonoids and phenolics have been found to have antilipidaemic properties^[64]^. The medicinal plant species Ziziphus lotus (L.) Lam. (Z. lotus fruit) [ZLF] is widely distributed across the Mediterranean basin and has a high flavonoid and phenolic content. The water-based extract of ZLF was shown to have strong anti-hyperlipidaemic and anti-atherogenic effects in a long-term high-fat diet model in albino mice^[64]^. It also showed significant antioxidant activity.

#### HDL-cholesterol and LDL-cholesterol

Malfunctions in the metabolism of HDL and LDL cholesterol have been linked to a higher risk of coronary artery disease, cerebrovascular accident, Alzheimer’s disease, myocardial infarction, and many other types of heart disease^[65]^. The rising incidence of Metabolic Syndrome and its correlation with low HDL cholesterol underscore its significance for diagnosis and treatment^[47]^. Studies of Africans have shown that lipoprotein particle sizes vary between ethnic groups. People of African descent have larger HDL and LDL particles and smaller VLDL particles, which makes their lipoprotein particles less likely to cause atherosclerosis^[66,67]^. Another investigation into interethnic heterogeneity in lipid profiles examined what this means for African Americans at risk of metabolic diseases who are not properly identified^[68]^. Blood lipids are clearly distributed differently across ethnic groups, with African Americans having a better serum lipid profile than other ethnic groups in the USA. These disparities may reflect differences in physical and genetic traits between ethnic groups, as shown among people of African heritage who today live in widely disparate contexts^[68]^. Nonetheless, a recent study in a Korean community revealed a favourable correlation between a raised TG/HDL-C ratio and a higher possibility of diabetes^[69]^

The current research showed that BCP substantially increased serum HDL (p**≤** 0.05) in animals with HFHFD after diet withdrawal, as well as between the withdrawn and continued diet groups. However, when HFHFD + BCP was compared to the corresponding continuing diet group, changes in LDL were not significant. These findings demonstrate how BCP affects HDL and LDL, helping to improve dyslipidaemia in cardiovascular dysfunction.

Research demonstrated that the BCP-containing aqueous and ethanolic extracts of A. squamosa leaves lowered cholesterol, triglycerides, and LDL cholesterol while raising HDL cholesterol^[70]^. The likely mechanism of action may involve initiating lipoprotein lipase activity and stimulating β-cells to release sufficient insulin to remove triglycerides from the plasma^[71, 72]^. Additionally, in a Spanish study relating Metabolic Syndrome to serum LDL, it was found that both low and high serum LDL levels were associated with a higher incidence of MS in working people, particularly among male employees. However, the frequency of MS in female workers was elevated only by high LDL-C levels (> 135.0 mg/dL) ^[73]^.

#### Triglycerides (TG) and Apo B

Studies have shown that low HDL cholesterol typically coexists with increased TG levels. This is because TG-rich lipid particles are cleared by HDL particles. As a result, HDL levels should be low when TG levels are high, and vice versa. “The dyslipidaemia of insulin resistance [IR]” is the common term used to describe the very atherogenic pattern of high TG and low HDL^[74]^. Apolipoprotein B (Apo-B), a protein mostly found in LDL-C, is used to predict the development of CVD without reference to other factors, particularly among individuals with type 2 diabetes (T2DM)^[75]^. Additional investigations have shown that ApoB and non-HDL-c are more strongly associated with coronary heart disease events than LDL-c, even in people who have MS ^[76]^. In addition, ApoB and non-HDL-c are higher in people with MS than in those without MS, regardless of LDL-c levels ^[76]^. It has been shown that ApoB and LDL-c are less in sync in people with MS or insulin resistance. ApoB seems a more reliable marker of CV risk than LDL-c ^[76]^. Another study also found that both traditional and new lipid profiles in Metabolic Syndrome showed increased atherogenicity^[76]^. Evidence showed that having LDL-c within goal was predicted by higher TG and HDL-c and lower ApoB ^[77]^. According to the same researchers, relying only on LDL-c may miss opportunities to lower CVD risk 60. Furthermore, ApoB, oxidised LDL-c, and especially non-HDL-c, may be useful indicators in assessing CVD risk in persons diagnosed with MS and may be therapeutic targets ^[77]^.

The current study results revealed a decrease in TG levels between the HFHFD continued+BCP and HFHFD withdrawn+BCP groups. This indicates that discontinuing the HFHFD diet in the presence of BCP led to a greater reduction in TG levels. However, APO B levels decreased in the BCP treatment groups, with the HFHFD withdrawn+BCP group showing lower levels than the HFHFD continued+BCP group. These findings demonstrate BCP’s modulatory effects on these markers of cardiovascular dysfunction and its ability to reduce metabolic dysfunction and cardiovascular damage caused by Metabolic Syndrome.

Comparing treated and untreated diabetic rats (control), previous studies have shown that the water-based extract of Annona squamosa substantially decreased triglyceride and total cholesterol levels while gradually increasing HDL cholesterol^[78]^. Hypertriglyceridaemia remains a primary risk factor for coronary artery disease and atherosclerosis, irrespective of other factors^[79]^. These disease entities constitute the primary global sources of morbidity and death. A key pathophysiological mechanism in hypertriglyceridaemia is impaired clearance of lipoproteins rich in plasma triglycerides. β-caryophyllene (BCP) has been shown to act as an agonist of the peroxisome proliferator-activated receptor (PPAR) isoforms (PPAR-α/γ) ^[79]^. BCP can be used to treat hypertriglyceridaemia in several ways^[80]^.

### Oxidative stress markers: Total protein (g/dl), GSH (Mm), SOD (u/ml), MDA (µm) and Nitric oxide (NO)

It is impossible to overstate the crucial role of oxidative stress (OS) in heart failure and Metabolic Syndrome. Research indicates that elevated OS in people with type 2 diabetes (T2DM) and Metabolic Syndrome is a major reason for their increased risk of heart disease^[61]^. OS is widely recognised as a biologically significant factor that promotes local inflammation and cardiovascular dysregulation by inducing vascular gene expression. Excessive reactive oxygen species (ROS) produced by vascular walls during OS damage endothelial cell integrity and function ^[81,82.]^

The current investigation evaluated oxidative stress indicators, namely: Malondialdehyde (MDA) (µM), Glutathione reductase (GSH) (mM), Superoxide dismutase (SOD) (u/ml), and Total protein (g/dl). According to research, excessive or insufficient synthesis of reactive oxygen species (ROS) and nitrogen species (RNS) leads to OS, which is linked to decreased antioxidant system quantity or expression, as well as reduced antioxidant activity^[81]^. ROS and RNS function as signal transduction molecules that support cellular activity and provide cellular protection when present in appropriately low concentrations. On the other hand, when produced in excess, as in inflammatory tissues^[81]^, they can generate more highly reactive molecules that irreversibly oxidise proteins, lipids, and nucleic acids. The oxidative modification of crucial enzymes or regulatory sites is particularly significant because it alters cell signalling and causes programmed cell death^[82]^. Experimental diabetic neuropathy is associated with oxidative stress, which results from increased free radical production and/or reduced antioxidant defences^[83]^.

Based on the current study results on the modulatory effects of Beta Caryophyllene on cardiovascular dysfunction indices in Metabolic Syndrome, different degrees of oxidative stress were observed across the diet/treatment groups. Reduced glutathione (GSH) levels in the Beta Caryophyllene treatment groups were substantially higher (p<0.05) in rats in which the HFFD diet was continued (BCP group, 46%) compared with the initial GSH levels in the HFHFD-only group. The GSH concentrations were also substantially raised in the HFHFD withdrawn + BCP group, in contrast to the HFHFD withdrawn only group. Animals treated with HFHFD + BCP showed no substantial change in total protein levels compared with the withdrawal diet group and the continuing diet group. SOD activity was significantly elevated in rats given Beta Caryophyllene upon discontinuing the HFHFD diet, although MDA levels had a substantial reduction (p<0.05) in those animals.

Furthermore, results from this study clearly show that, across various response levels in specific diet/treatment groups, high-calorie diets increase oxidative stress. The data also indicate that oxidative stress is ameliorated after treatment with Beta-caryophyllene, as evidenced by differential outcomes. Beta-caryophyllene treatment was shown to ameliorate oxidative stress. Beta-caryophyllene could serve as a possible novel future management strategy for Metabolic Syndrome. Hence, this study demonstrated Beta-caryophyllene as a natural antioxidant, with the best potency when administered after withdrawal from the Metabolic Syndrome diet.

Similarly, a study on spinach (rich in BCP) that examined how consuming spinach and engaging in aerobic exercise (AE) could improve abnormalities linked to Metabolic Syndrome in rats revealed astounding results regarding oxidative stress status^[84.]^ Rats fed fructose had considerably (p < 0.01) higher MDA levels, a lipid peroxidation marker, in their hearts than those in the normal group. Elevated MDA output was markedly (p < 0.01) inhibited by all treatments, while rats in the fructose control group showed a substantial decrease (p < 0.01) in concentrations of GSH and SOD, and in the activities of CAT, GPx, and GR in heart homogenates compared with the normal set^[84]^. NAOE, AE, and gemfibrozil treatments restored these cardiac antioxidants, although AE didn’t restore the low GR levels^[84]^. [Spinacia oleracea NAO-rich extract (NAOE); Reduced glutathione (GSH); Superoxide dismutase (SOD); Catalase (CAT); Glutathione reductase (GR); Glutathione peroxidase (GPX); Natural antioxidants (NAO); Aerobic exercise (AE)]^[84]^.

It is not straightforward to test the antioxidant outcome of every quantity for each item used in a trial on the balance of natural antioxidants that protect against Metabolic Syndrome by lowering oxidative stress^[85]^. However, to find the best mix of natural antioxidants, such as vitamin C, phenols from green tea, and proanthocyanidin preparations from grape seeds, experts used the Hill equation to create “The Fixed Dose Combination (FDC)” mathematical model^[71]^. The experts looked at how FDC affected OS, blood sugar, and lipid levels in adipocytes (3T3-L1) grown in the lab. The experiment utilised HFD-fed rats and KK-ay mice as obesity and diabetes models, respectively. The FDC had strong antioxidant and anti-glycation effects, lowering lipid peroxidation (reducing MDA levels)^[85]^. These experts say that FDC might be a good way to treat people with diabetes and obesity, as well as a way to lose weight and make insulin work better. They said this mathematical method could be a fresh and useful way to think about how natural antioxidants can be combined, and it might help create new medicines that could help people worldwide with metabolic diseases^[85]^.

In animal models of type 2 diabetes mellitus (NIDDM), a water-based Annona squamosa leaf solution, rich in beta-caryophyllene, altered the lipid profile and antioxidant enzymes^[70]^. Malondialdehyde levels declined significantly across all tissues when the enzymes catalase (CAT), superoxide dismutase (SOD), reduced glutathione (GSH), glutathione reductase (GR), and GST performed better^[83]^. Beta-caryophyllene (BCP) is a potent antioxidant and free radical scavenger that can neutralise unstable free radicals such as superoxide and hydroxyl anions^[84]^. BCP also normalises the glutathione redox cycle^[85]^. BCP may inhibit lipid degradation. A recent study found that BCP prevented hypoxic neuroinflammatory processes and damage^[73]^. It achieved this by reducing mitochondrial ROS generation, activating NF-κB in microglia, and preventing the release of pro-inflammatory cytokines^[85]^. Additionally, studies have shown that BCP is superior to several common antioxidants, including the therapeutically utilised vitamins C and E ^[83]^, and it regulates a number of genes related to oxidative stress, antioxidant defence, xenobiotic detoxification, ageing, and lifespan^[75]^.

Recent studies suggest that body fat dysfunction and insulin resistance are promoted by depletion of nitrate and nitrite, persistent end-products of nitric oxide (NO)^[86]^. Furthermore, research has already shown that the primary causes of endothelial dysfunction in Metabolic Syndrome are decreased endothelial nitric oxide synthase (NOS) activity and NO bioavailability^[86]^. Notably, research has shown a connection between the leptin signalling system and the nitric oxide (NO) pathway, since leptin stimulates cardiomyocytes to produce NO^[87–94]^.

This study revealed that the HFHFD-withdrawn + BCP group had lower serum nitric oxide levels than the HFHFD-continued + BCP group. However, substantially higher nitric oxide levels were observed in the HF/HFD-continued + BCP group compared with the HF/HFD-continued group, and in the HFHFD-withdrawn + BCP group compared with the HF/HFD-withdrawn group. These results describe the modulatory effects of BCP on nitric oxide levels, showing amelioration of cardiovascular dysfunction in metabolic syndrome.

Other research has shown that a range of medicinal plant extracts may alleviate MS disease ^[12, 88]^. The aqueous extract of Piper sarmentosum, which has been shown to be rich in BCP, increased the production and activity of eNOS in the thoracic aorta, raised serum eNOS levels, and increased NO in the mesenteric artery in rats with hypertension induced by N ω-nitro-L-arginine methyl ester hydrochloride (L-NAME)^[95]^. Moreover, NO is a strong vasodilator^[89]^. Research has also shown strong anti-inflammatory benefits of NO, but just as many studies show that NO may worsen tissue and cellular failure caused by inflammation^[95]^. Understanding the metabolic chemistry of NO and its by-products will provide a framework for distinguishing between the negative and pro-inflammatory effects of NO and its positive and anti-inflammatory effects, even though the precise causes of these seemingly contradictory observations are unknown^[95]^.

Nitric oxide has both direct and secondary activities^[95.]^ NO-derived intermediates, such as reactive nitrogen oxide species formed from NO-oxygen or superoxide reactions, induce indirect effects. NO interacts directly with biological molecules or targets to produce direct effects. These specialists argue that separating NO chemistry into these two categories enables chemical processes to be classified by NO production rate. For instance, low NO fluxes may have direct impacts, whereas high NO fluxes may have secondary consequences. Since constitutive isoforms of nitric oxide synthase (NOS), such as brain NOS or those in vascular cells, generate minimal NO, NO’s direct effects should be greater. Under normal conditions, cells produce modest but significant quantities of NO and reactive oxygen species, so direct NO chemistry is most prevalent in healthy tissue. In tissues with chronic inflammation and high-output inducible NOS (iNOS) levels, nitrosation, nitration, and oxidation occur more commonly. These researchers^[95]^ claim that understanding the location, timing, and rates of NO generation helps identify the relevant targets and the type of chemistry most likely to be affected.

Numerous facets of the heart’s excitation-contraction coupling and energy metabolism are affected by NO synthase (NOS) isoforms, which localise to distinct subcellular microdomains^[96]^. NOS1 localises to the cardiac SR under physiological conditions, where it probably contributes to the S-nitrosylation of the anti-goat ryanodine receptor (RYR), thereby facilitating Ca2+ release^[97]^. A study found that decreased neuronal NOS expression contributes to oxidant stress and nitroso-redox mismatch in the hearts of ob/ob mice^[98]^. In an obesity mouse model, leptin insufficiency is linked to decreased NOS1 levels in the heart without relocation, resulting in nitroso-redox imbalance and the emergence of heart failure-related symptoms^[99]^. Therefore, the researchers hypothesised that nitroso-redox imbalance associated with NOS1 represents a universal mechanism for cardiac dysfunction that extends beyond overt heart failure^[99]^.

Numerous in vitro and in vivo studies have shown that β-caryophyllene (BCP) has potent anti-inflammatory properties^[15]^. One of its most well-established effects is modulating cytokines and chemokines that drive inflammation, thereby preventing the initiation and progression of immune-inflammatory diseases via CB2 receptors^[73,75]^. Numerous studies have demonstrated that CB2 receptor stimulation enhances BCP’s anti-inflammatory activity. After binding to CB2 receptors, BCP inhibits adenylate cyclase, induces Ca2+ changes within cells, and further activates the signalling pathways controlled by p38 and Erk1/2^[15]^. Additionally, high concentrations of BCP have been linked to anti-inflammatory properties in several plants, including Copaifera multijuga^[100]^.

#### Organ integrity

The chronic injurious diet specifically caused infarction of cardiac tissue. Pathology in other organs on the chronic Ms diet included fibrosis, fat cell deposits, and inflammatory cells. BCP reduced inflammation, fat cell deposits, and fibrosis, thereby improving overall organ architecture. BCP administration also ameliorated heart enlargement and hypertrophy caused by a chronically induced high-fat, high-fructose diet. The metabolic syndrome diet affected cardiac tissue, making it heavier, as evidenced by increased organ weight. The architecture also became disorganised. Myocardial infarcts were observed as a result of the inflammatory process, which affected the heart muscle (intima, media, and externa). The myocardium showed abnormal myocytes. However, BCP reversed these processes, restoring a healthy heart upon administration. BCP ameliorated liver injury among the rats fed on the dangerous high-calorie diet in this study, as shown by a marked decrease in steatosis and fibrosis in liver architecture^[101]^, as previously explained. BCP also ameliorated liver injury by decreasing hepatomegaly and macrovesicular steatosis.

### Public health and translational implications of Beta-caryophyllene intervention

The findings of this study demonstrate that Beta-caryophyllene (BCP) significantly restores hemodynamic metrics like blood pressure and heart rate while concurrently improving critical metabolic and inflammatory biomarkers in a diet-induced rat model of metabolic syndrome. Translating these preclinical results into human population health frameworks is highly relevant, given that metabolic syndrome and its associated cardiovascular dysfunctions represent an escalating global non-communicable disease (NCD) crisis. Lifestyle and dietary shifts, characterized by the heavy consumption of ultra-processed foods rich in fats and high-fructose corn syrup, mirror the experimental induction diet used in this study and drive the cardiovascular mortality burden worldwide. Traditional pharmaceutical interventions for cardiometabolic disorders are frequently constrained by high financial costs, systemic adverse effects, and poor accessibility—particularly in low- and middle-income countries (LMICs) such as Nigeria, where NCDs account for over 30% of all mortalities. Consequently, validating low-cost, safe, and scalable dietary nutraceuticals like BCP offers a practical paradigm shift toward sustainable primary prevention strategies at the population level.

From a clinical and molecular standpoint, the multi-target efficacy of BCP observed in this study provides a strong rationale for its development as a human functional food component. In male Wistar rats, BCP effectively counteracted the pathological hallmarks of insulin resistance, showing robust anti-atherogenic, anti-inflammatory, and antioxidant actions evidenced by the down-regulation of apolipoprotein-B, systemic C-reactive protein (CRP), and lipid peroxidation. In humans, elevated Apo-B and high-sensitivity CRP are powerful, independent predictors of atherothrombosis and future myocardial events. Because BCP is a naturally occurring dietary cannabinoid abundantly found in common, culturally accepted spices like black pepper (*Piper nigrum*), cloves, and rosemary, it possesses an inherent safety profile recognized as Generally Recognized as Safe (GRAS) by international regulatory agencies. This status vastly simplifies the translational pathway, allowing BCP-rich botanicals or fortified functional ingredients to be seamlessly integrated into national nutrition policies, public health education programs, and targeted dietary interventions aimed at managing metabolic syndrome before it progresses to overt, irreversible heart failure.

However, moving from these encouraging animal data to practical public health applications requires a careful, phased translational framework. While the 50 mg/kg oral dose in rats demonstrated clear cardioprotective benefits and successfully reversed tissue microarchitecture distortion in vital organs, human equivalent dosing (HED) must be rigorously established through pharmacokinetic scaling. Future human clinical trials must also evaluate the long-term bioavailability, metabolic clearance, and optimal delivery matrices of BCP, as sesquiterpenes can vary in stable systemic absorption. Additionally, public health advocacy should focus on protecting and promoting regional agricultural systems that cultivate these bioactive-rich food sources. Aligning natural product research with the World Health Organization’s Global Action Plan for the Prevention and Control of NCDs provides a structural roadmap to curb chronic disease burdens, directly supporting Sustainable Development Goal (SDG) 3.4 to reduce premature mortality from NCDs by one-third by 2030.

### Research highlights

High-fat high-fructose diets successfully induce severe hemodynamic and electrophysiological abnormalities in Wistar rats.

1. Beta-caryophyllene treatment significantly restores systolic blood pressure, heart rate, and prolonged PR intervals.
2. Oral administration of beta-caryophyllene decreases atherogenic apolipoprotein-B, total cholesterol, and systemic C-reactive protein.
3. Myocardial tissue injury, lipid peroxidation, and macrovesicular steatosis are prominently reversed by the sesquiterpene.
4. The natural cannabinoid offers an affordable, GRAS-designated nutraceutical strategy for global public health NCD prevention.

### Conclusion

The mammalian heart is a vital organ in human physiology and must be maintained in optimal condition. The influence of diet on its maintenance cannot be overstated. This study demonstrated, via serial physiological measurements that served as markers of cardiovascular function, that Beta-caryophyllene is a potential therapy for cardiovascular dysfunction in Metabolic Syndrome. It also showed that beta-caryophyllene (BCP) is cardioprotective. The study further showed that withdrawing an HFHFD diet excessively rich in calories may improve the outcome of a metabolic disease in progress. Therefore, it is reasonable to conclude that a diet rich in BCP, via fruits and vegetables, alleviates and prevents cardiovascular dysfunction. This represents a proposed therapeutic approach to managing cardiovascular morbidity in Metabolic Syndrome. Beta-caryophyllene ameliorated Metabolic Syndrome-induced cardiovascular dysfunction in male Wistar rats through its anti-atherogenic and antioxidant properties. Given that BCP is found in common spices, food and plants, as discussed, and is GRAS, dietary incorporation could be a low-cost public health strategy for NCD prevention in Africa.

## Acknowledgments

The authors acknowledge the following Departments:

Physiology, Biochemistry, Pharmacognosy & Herbal Medicine, and Veterinary Medicine, all of the University of Ibadan, Nigeria.

## Supporting information

**S1 Table. Modulatory Outcome of BCP treatment on Heart rate and Blood Pressure (BP) Variables among HFHFD animal sets on HFHD diet**.

**S2 Table. Modulatory outcome of Beta Caryophyllene (BCP) on ECG variables among HFHFD-Rats on HFHFD**

**S1 - S11 Figs. Modulatory outcomes of BCP on CRP, APO B, lipid profile and oxidative stress respectively.** S1-S11 Legend:

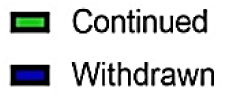

**Values in Figs 3-6** were presented as means and standard deviations for 12 animals. *= statistically significantly different from HFHFD-continued, **= statistically significantly different from HFHFD-withdrawn. S3-S6 Legend: HF/HFD = High-fat and High-fructose diet, BCP = Beta Caryophyllene.

**Values in Figs 7-11** were presented by means and standard deviation of 12 animals. *= statistically significantly different from HF/HFD-continued, **= statistically significantly different from HFHFD-withdrawn. S7-S11 legend: HF/HFD= High fat and High fructose diet, BCP= Beta Caryophyllene

**Figs 12-14 (Plates A-E). Photomicrographs of the modulatory outcomes of BCP on organ tissues (cardiac and hepatic tissues)**

## PHOTOMICROGRAPHS

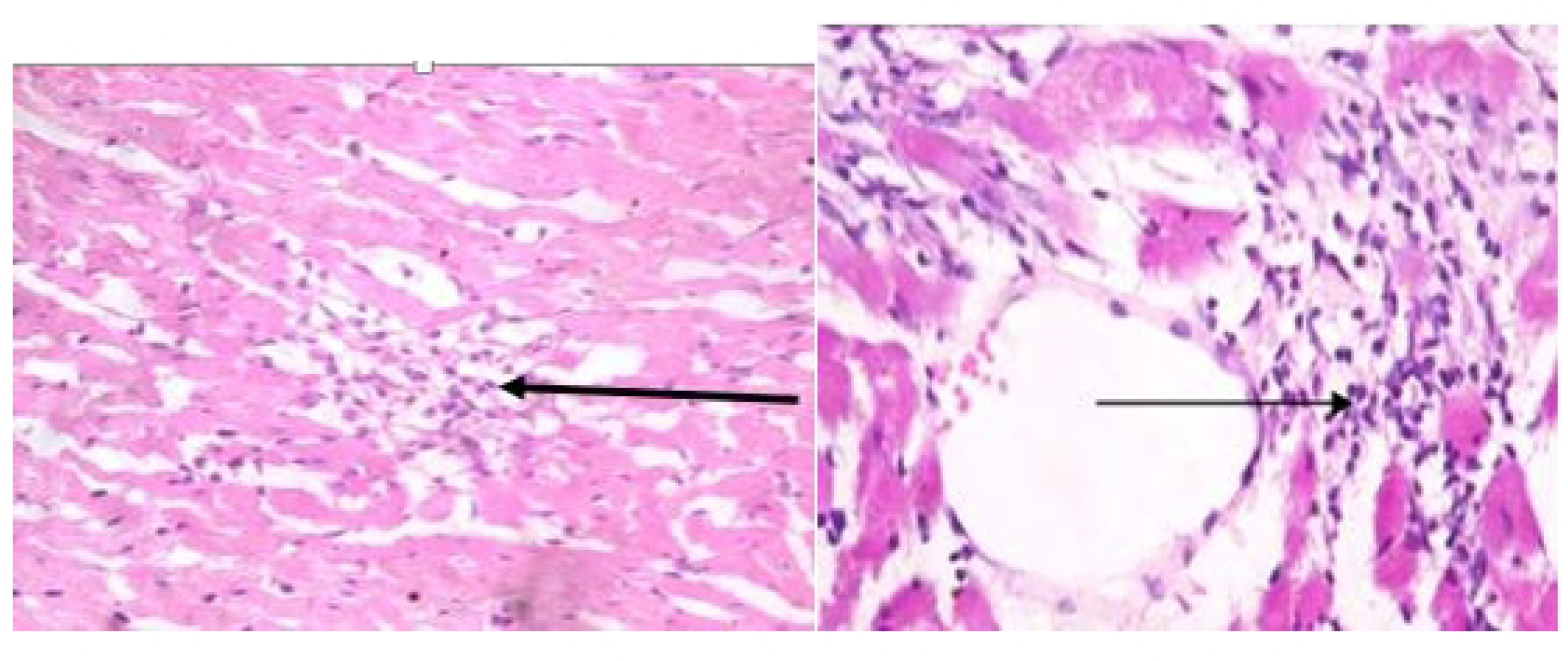

Plate A (left): HFHFD + BCP Cardiac tissue photomicrographs reveal a slight degree of inflammation (black arrow).

Plate B (right): HFHFD without BCP Cardiac Tissue Photomicrographs show moderate infarction (black arrow) and moderate infiltration by inflammatory cells (thin arrow). The infarcted area is sharply demarcated from relatively preserved myocardium i.e, heart muscle is preserved.

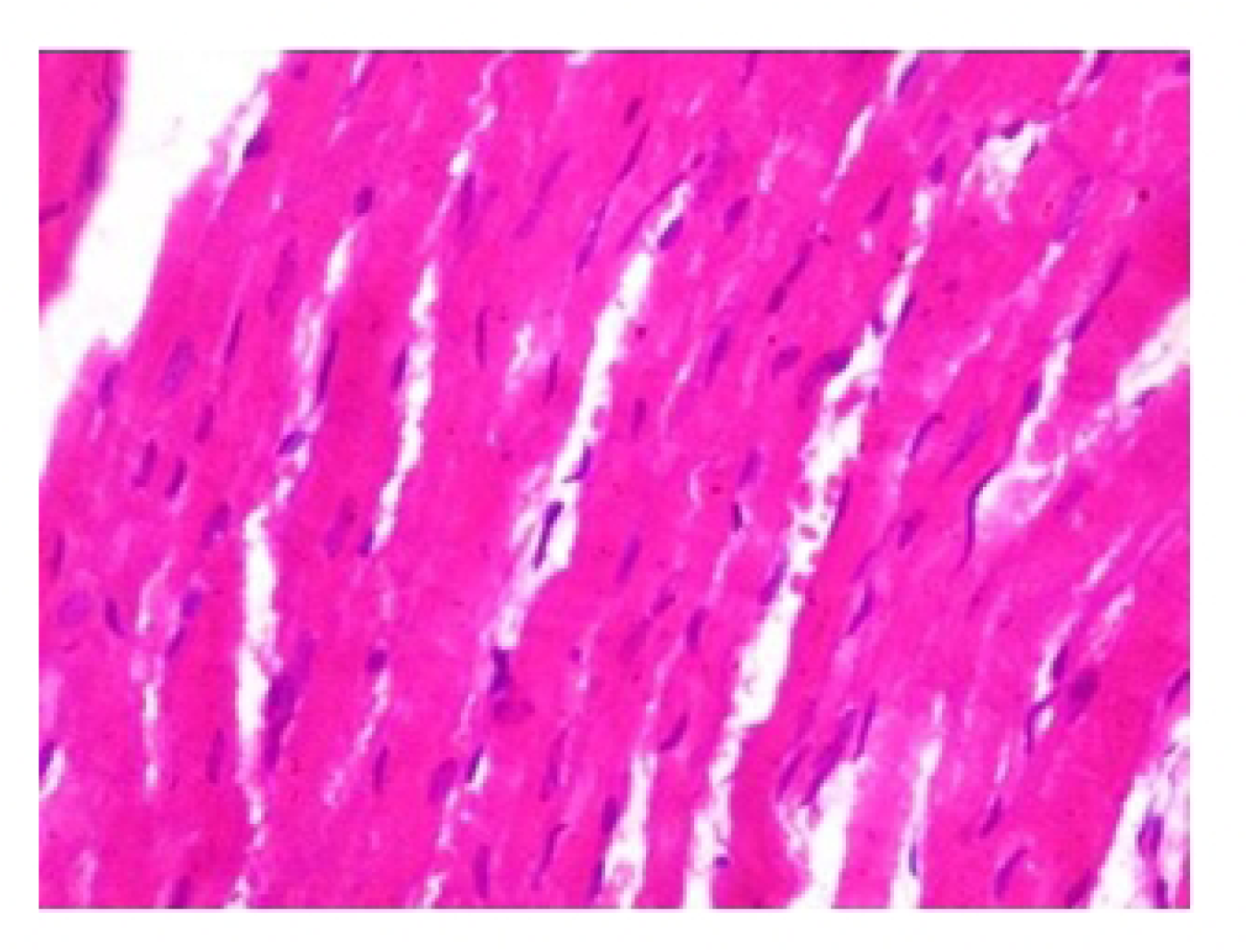

Plate C: High fat + high fructose diet (withdrawn) + Bela Caryophyllene Cardiac tissue photomicrographs show normal architecture and cellularity of myocytes. No significant lesion seen. Intact heart muscle without inflammation.

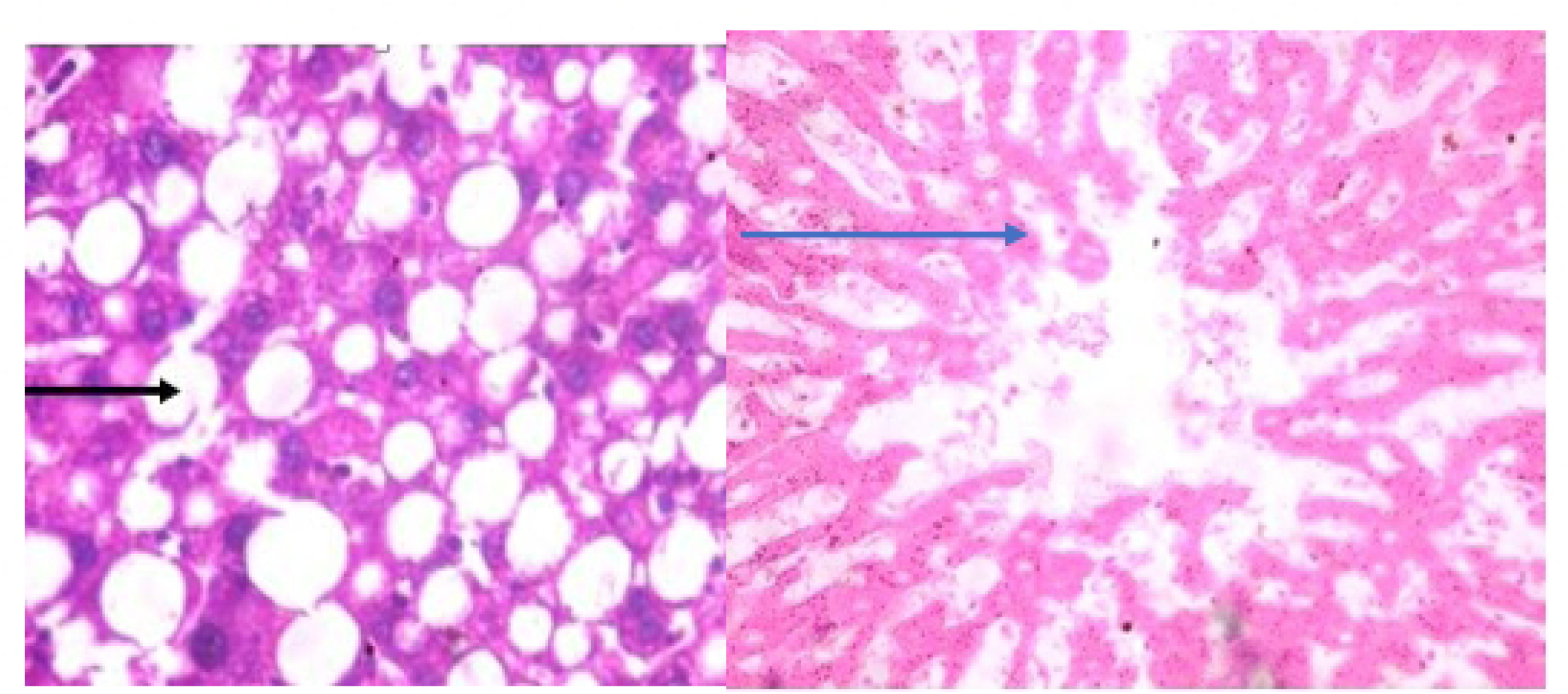

Plate D (left): High fat+ high fructose diet (continued) + Beta caryophyllene hepatic tissue photomicrographs reveal widespread, severe macro vesicular steatosis (black arrow).

Plate E (right): High fat + high fructose diet (withdrawn) + Beta caryophyllene hepatic tissue photomicrographs display sinusoids, portal triads, central veins, and healthy hepatocytes. No notable lesion was observed.

## Notes

### Competing Interest Statement

The authors have declared no competing interest.

